# Apparent Diffusion Coefficient fMRI shines light on white matter resting-state connectivity compared to BOLD

**DOI:** 10.1101/2024.07.03.601842

**Authors:** Inès de Riedmatten, Arthur P C Spencer, Wiktor Olszowy, Ileana O Jelescu

## Abstract

Resting-state functional magnetic resonance imaging (fMRI) is used to derive functional connectivity (FC) between brain regions. Typically, blood oxygen level-dependent (BOLD) contrast is used. However, BOLD’s reliance on neurovascular coupling poses challenges in reflecting brain activity accurately, leading to reduced sensitivity in white matter (WM). WM BOLD signals have long been considered physiological noise, although recent evidence shows that both stimulus-evoked and resting-state WM BOLD signals resemble those in gray matter (GM), albeit smaller in amplitude. We introduce apparent diffusion coefficient fMRI (ADC-fMRI) as a promising functional contrast for GM and WM FC, capturing activity-driven neuromorphological fluctuations. Our study compares BOLD-fMRI and ADC-fMRI FC in GM and WM, showing that ADC-fMRI mirrors BOLD-fMRI connectivity in GM, while capturing more robust FC in WM. ADC-fMRI displays higher average clustering and average node strength in WM, and higher inter-subject similarity, compared to BOLD. Taken together, this suggests that ADC-fMRI is a reliable tool for exploring FC that incorporates gray and white matter nodes in a novel way.

## Introduction

First introduced in 1995 by Biswal et al. ^1^, resting-state functional MRI (rs-fMRI) historically studies the spontaneous low frequency oscillations (*<* 0.1 Hz) of the Blood Oxygen Level-Dependent (BOLD) signal at rest^2,3^. The coherent functional patterns observed consistently across individuals are referred to as the Resting State Networks (RSNs), such as the Default Mode Network (DMN), originally defined by Raichle et al. ^4^ using PET. Typically, brain functional connectivity (FC) can be built from temporal statistical dependence between MRI time series across different brain regions and summarized as networks^5^. In this context, an anatomical or functional parcellation forms the nodes of the network, while a measure of FC forms the network edges (“weights” or “connections”)^6^. Subsequently, the properties of FC networks can be evaluated by computing graph theoretical metrics on brain networks. The ability of FC to capture significant changes in normal resting-state brain activity has clinical relevance in the context of disease or disorder^7^. For instance, one promising application of rs-fMRI is as a potential aid for preoperative surgical planning to localize the eloquent cortex, as proposed by Zhang et al. ^8^ for brain tumor resection. Stampacchia et al. ^9^ also provided preliminary evidence that rs-fMRI could be used for clinical fingerprinting, to detect individual FC alterations following cognitive decline.

Most of the fMRI studies to date rely on the BOLD contrast, which is a very informative yet indirect measure of brain activity (in this paper, “brain activity” is understood as both neural processing in the GM and action potential propagation in the WM). Indeed, the BOLD signal originates from neighboring blood vessels, not directly from neural activity, thus its area extends beyond the site of the active neurons. In addition, the BOLD response also has limited temporal specificity, as its onset is delayed relative to the brain activity onset. Furthermore, it displays reduced sensitivity in the WM regions, where BOLD low frequency oscillations, blood flow and blood volume are reduced by 60%^1,10^, 75% and 75%, respectively^11^. Finally, when vasculature is impaired, such as observed in healthy aging^12^, brain tumors^13^, multiple sclerosis^14^ or using pharmacological agents such as antihistaminic or anesthesic drugs^15^, the BOLD response can be altered^16^, even when the brain activity is not affected^17^.

As a result of limited sensitivity, BOLD-fMRI signal in the WM has historically been regressed out as a nuisance variable, although there is now compelling evidence for the contribution of neural activity to WM BOLD signals, in both stimulus-evoked and resting-state settings ^11,18,19^. Indeed, Li et al. ^20^ derived BOLD hemodynamic response functions (HRFs) in the superficial, middle and deep WM using the Stroop test, supporting a stimulus-induced BOLD signal change in the WM. In addition, voxel-wise WM activity maps were derived using a visual stimulus^21^. Although the GM-GM connectivity is the most widely studied connectivity in rs-fMRI, the GM-WM^22,23^ as well as WM-WM connectivity^21,24^ have also gathered recent attention, revealing reproducible GM-WM and WM-WM resting-state networks.

The potential of using the apparent diffusion coefficient (ADC) of water molecules as an alternative rs-fMRI contrast was first investigated in 2020 in mice^16^ and in 2021 in humans^25^. As highlighted by Darquíe et al. ^26^, this diffusion-based contrast is sensitive to brain activity via neuromorphological or neuromechanical coupling, although the precise mechanism behind ADC-fMRI remains incompletely understood^16^. Briefly, fluctuations in tissue microstructure resulting from brain activity influence the random Brownian motion of water molecules, and are detected as a transient alteration of the ADC at the voxel-level^27^. A decrease in ADC was thus observed immediately after the application of a stimulus^25,28^, possibly due to neuronal swelling. Indeed, nanometer-scale swelling of neuronal somas^29^, neurites^29^, myelinated axons^30^, synaptic boutons^31^ and astrocytic processes^32^ were observed during brain activity. The cellular deformations assumed to induce a change in ADC have been shown with optical techniques. In terms of timescale, axonal displacement could be observed for 1 s optogenetic stimuli in mouse brain slices^30^. It started shortly (*<* 250 ms) after the stimulation onset, and lasted 1-2 s after the stimulation stopped. For GM, Ling et al. ^29^ observed spike-induced single-cell neuronal deformation using high-speed interferometric imaging, and showed that the deformation started with a latency of less than 0.1 ms latency, and lasted about 2 ms for a single action potential. A train of action potentials would be expected to maintain the deformation for the duration of the train, similar to the axon displacement measured during sustained optogenetic stimulation. Furthermore, it cannot be ruled out that astrocytic deformations concurrent with brain activity also contribute to the ADC-fMRI contrast, for which timings may be delayed^33^.

Since it was first proposed by Darquíe et al. ^26^, diffusion fMRI (dfMRI) has faced criticism, including potential BOLD contamination^34^ and limited sensitivity^35^. The non-vascular nature of the diffusion-weighted functional signal has however been extensively documented in the literature. It was shown in living mice that aquaporin-4 blocker (astrocyte function perturbator) notably enhanced the BOLD response induced by optogenetic activation of astrocytes, while showing no impact on the dfMRI signal^36^. Similarly, when rats were exposed to the neurovascular coupling inhibitor nitroprusside (causing vasodilation and hypotension), the BOLD signal was notably affected, while the dfMRI signal was not^37^. These findings suggest an underlying mechanism for dfMRI which is independent of neurovascular coupling. Nonetheless, under hypercapnia, which should elicit vasodilation without changes in neural activity, the diffusion-weighted fMRI signal up to b = 2400 s mm^−2^ exhibited BOLD-like fluctuations^34^. Indeed, the diffusion-weighted signal demonstrates sensitivity to *T*_2_ variations 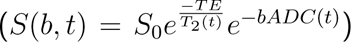, essentially capturing the fundamental mechanism underlying BOLD signal. ADC-fMRI can be used to overcome this limitation by calculating an ADC time series from the diffusion-weighted signals. The ADC is calculated from a ratio of two diffusion-weighted signals, where the *T*_2_-weighting is largely cancelled out (Eq. 1).

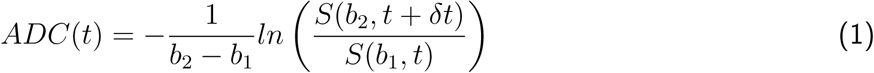

However, *T*_2_-weighting is not the sole vascular contribution to the dfMRI signal that requires mitigation. During the hemodynamic response, magnetic susceptibility-induced background field gradients (G*_s_*) vary around blood vessels^38^. These gradients impact the diffusion-weighted signal and the ADC by contributing to the effective b-value (Eq. 2).

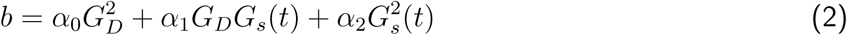

To counteract this, acquisition schemes which minimize cross-terms between diffusion gradients and background gradients, such as Twice-Refocused Spin-Echo (TRSE), can be used instead of the traditional Pulse-Gradient Spin-Echo (PGSE) sequence^38^.

Finally, the selection of the two b-values b_1_ and b_2_ for ADC calculation is also crucial. In previous studies, b_1_ was often set to 0. However, applying a small diffusion weighting instead (b_1_ = 200 s mm^−2^) suppresses most of the contribution from fast moving spins in the blood water pool and is therefore recommended to minimize the sensitivity of ADC functional changes to blood volume changes in the voxel^25,39^. Indeed, diffusion mediated signals obtained using b-values in the range 0-250 s mm^−2^ also include blood flow in capillaries (perfusion, also referred to as pseudo-diffusion in intra-voxel incoherent motion contrast). When using b_1_ = 200 s mm^−2^ as a cut-off, capillary perfusion on average only accounts for 1% of the measured diffusion signal^40^ and is considered negligible. We set b_2_ to 1000 s mm^−2^ as a compromise between diffusion mediated response amplitude vs SNR, as well as staying in the Gaussian Phase Approximation regime. Using higher b-values in the ADC calculation (in this case, b_1_ = 200 and b_2_ = 1000 s mm^−2^) is also referred to as “synthetic ADC”^41^ or “shifted ADC” (sADC)^33,42^.

Taken together, these criteria provide an ADC-fMRI acquisition that is closely related to brain activity, both spatially and temporally, relying on neuromorphological coupling rather than neurovascular coupling. Furthermore, ADC-fMRI by design should be much less affected by physiological noise that fluctuates on slower timescales than the TR. Indeed, cardiac and respiratory fluctuations are expected to be largely canceled out when taking the signal ratio of two consecutive images to calculate an ADC (Eq. 1), and ADC-fMRI thus to generate more reproducible results^10^. Finally, ADC-fMRI could be particularly beneficial in the WM, where BOLD-fMRI sensitivity is dramatically reduced^1,11^. As microstructural changes during brain activity also occur at the axonal level^30^, the applicability of resting-state ADC-fMRI can be extended to WM activity, as already shown using task-fMRI^43^.

The aim of this study was to examine the feasibility of utilizing ADC timecourses in rs-fMRI and to determine whether it could enable the exploration of GM-WM and WM-WM functional connectivity better than BOLD-fMRI. While BOLD signal is known to be stronger at higher field strength, the ADC signal is not expected to be affected by field strength. However, higher field strength typically results in better SNR, possibly impacting FC. Therefore, resting-state ADC-fMRI and BOLD-fMRI were acquired at 3T and 7T. We derived FC matrices for ADC-fMRI and BOLD-fMRI between GM as well as WM region-of-interests (ROIs). We studied the FC in three connectivity domains (GM-GM, WM-WM and GM-WM). We then used graph analysis to study the differences between the ADC-fMRI and BOLD-fMRI networks in terms of integration and segregation, and performed an inter-subject similarity analysis to assess the consistency of FC patterns across subjects for each contrast. This study confirms that ADC-fMRI mirrors the positive correlations observed in BOLD-fMRI. While comparable average clustering and average node strength were found for GM-GM connections, a higher average clustering and average node strength for ADC-fMRI in WM-WM edges suggest that it captures different FC to BOLD in the WM. In addition, a significantly higher FC similarity between subjects for ADC-fMRI than BOLD-fMRI in WM-WM connections suggests a higher reliability of ADC-fMRI in this brain tissue type, demonstrating its broader applicability across the entire brain, possibly due to reduced sensitivity to physiological noise. Taken together, these results encourage the use of ADC-fMRI, together with careful mitigation of vascular contributions, to further investigate WM functional connectivity.

## Results

The analysis was performed on ADC-fMRI and BOLD-fMRI data acquired at 3T (n*_ADC_*_−_*_fMRI_* = 12, n*_BOLD_*_−_*_fMRI_* = 10), and 7T (n*_ADC_*_−_*_fMRI_* = 10, n*_BOLD_*_−_*_fMRI_* = 10), separately. ADC-fMRI was derived from alternating b-values of 200 and 1000 s mm^−1^ acquired with TRSE echo-planar imaging (EPI) sequence, following Eq. 1. BOLD-fMRI was acquired with SE-EPI. As both field strengths led to similar results and interpretations, only the 3T results are discussed here while the 7T results are shown in the Supplementary Fig. 1-5. As the sampling period of BOLD-fMRI (1 s) is half the sampling period of ADC-fMRI (2 s), the results are also derived after downsampling BOLD-fMRI to the same sampling rate as ADC-fMRI (see Supplementary Fig. 6-10). Unless otherwise stated, BOLD-fMRI results are similar with a sampling period of 1 s or 2 s.

### Functional connectivity

FC matrices were derived for ADC-fMRI and BOLD-fMRI with Pearson’s partial correlation between each pair of atlas ROI average timecourses, regressing out the the average timecourse of the brain (Fig. 1A). The FC matrices are transformed to z scores using Fisher transform. Significant positive and negative FC weights within each contrast were determined with two-sided one-sample Wilcoxon tests with multiple comparisons correction, and split according to their connectivity domains (GM-GM, WM-WM, GM-WM).

**Figure 1:**
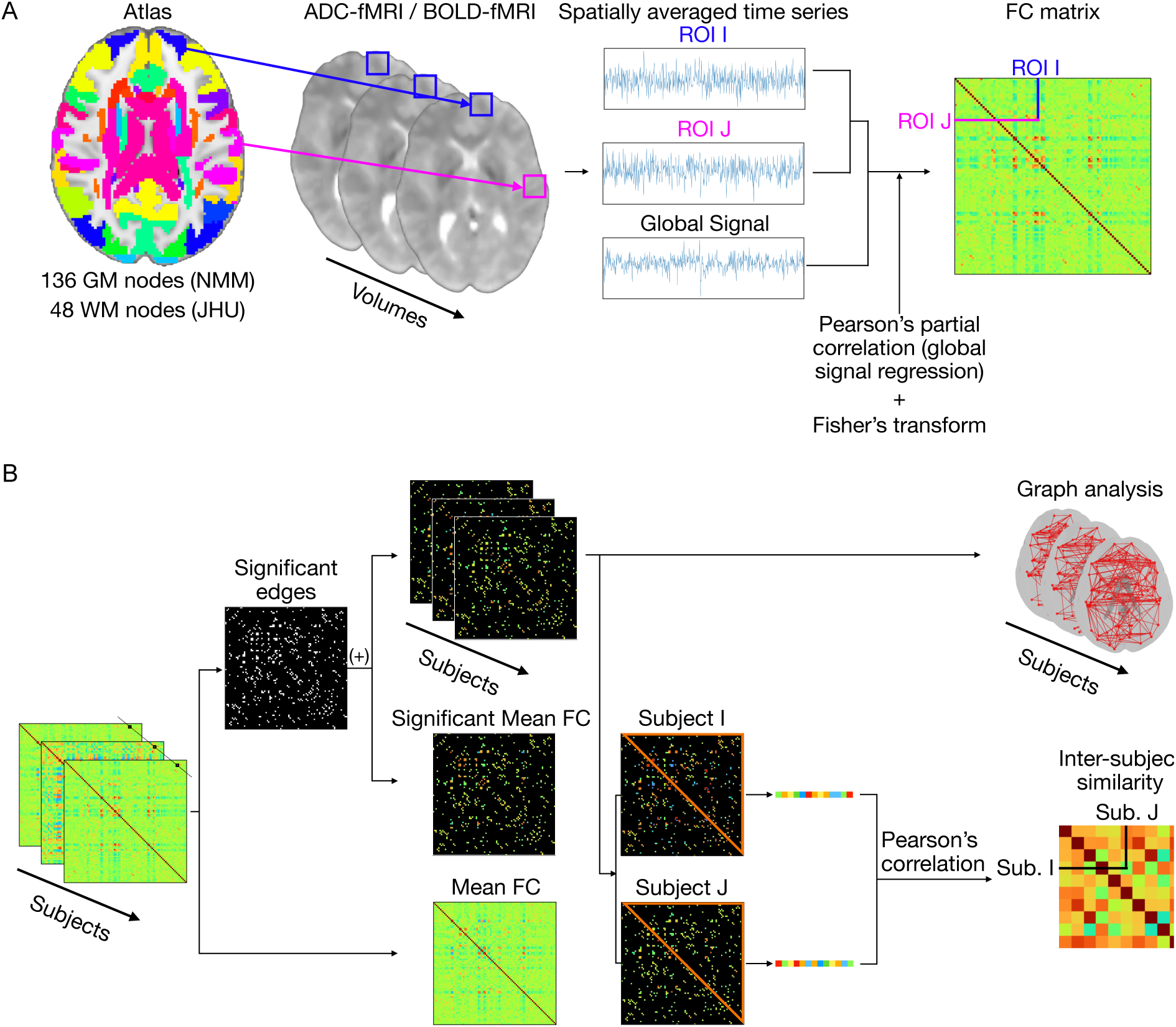
Methods for processing and analyzing the ADC-fMRI and BOLD-fMRI data. A) Subject-wise processing. Time series were averaged within ROIs shared in all subjects and containing at least 10 voxels. The correlation between a pair of ROIs was calculated using Pearson’s pairwise partial correlation with the global signal from the whole brain slab as a covariate. A Fisher transform was then applied to obtain the function connectivity matrix. B) Group-wise analysis. The binary mask of significant edges was derived using two-sided one-sample Wilcoxon tests with multiple comparisons correction (FDR Benjamini-Hochberg). It was then applied to individual FC matrices or to the mean FC matrix. The masked individual FC matrices were either transformed into graphs, on which graph metrics were calculated, or used to derive an inter-subject similarity index. For pairs of subjects, the edges contained in the upper triangle (excluding self-connections) were linearized, and a pairwise Pearson’s correlation coefficient was calculated.

### GM-GM connectivity

At 3T, FC matrices (Supplementary Fig. 11) yielded similar positive correlations between ADC-fMRI and BOLD-fMRI, with BOLD-fMRI displaying overall larger mean positive correlations in the GM-GM connectivity. Regions corresponding to the DMN were typically positively correlated to each other for both ADC-fMRI and BOLD-fMRI (Fig. 2, green edges, and Supplementary Fig. 11). The main regions of the DMN include the medial prefrontal cortex (the superior frontal gyrus medial segment -MSFG-, the medial frontal cortex -MFC-, the frontal pole -FRP- and the anterior cingulate gyrus -ACgG- here), the posterior cingulate (PCgG) and precuneus (PCu) and the lateral parietal cortex (the superior parietal lobule here)^4^. We found that the ACgG was positively correlated with the MFC and the MSFG, for both BOLD-fMRI and ADC-fMRI.

**Figure 2:**
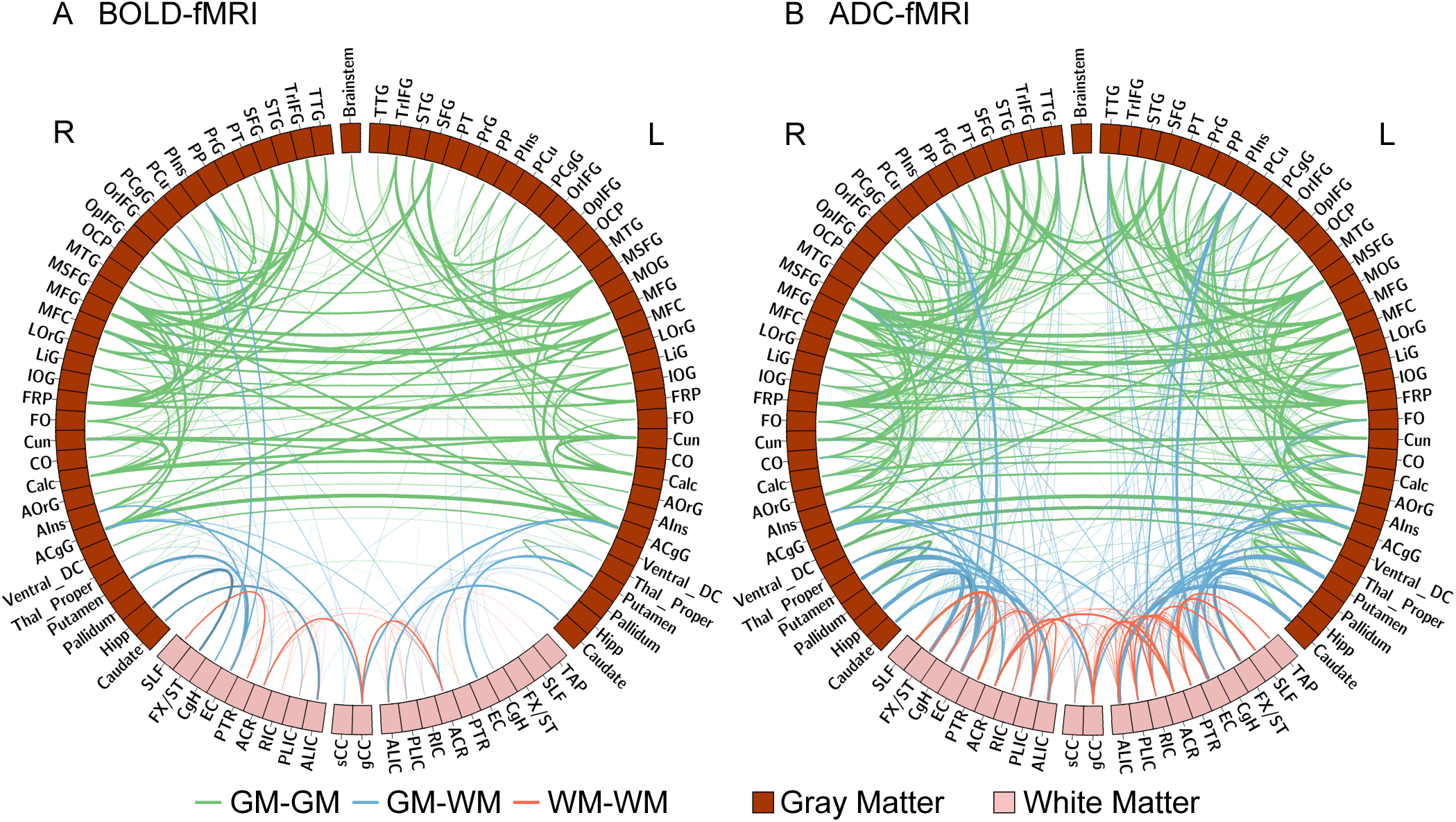
Mean FC of significant positive edges. A) BOLD-fMRI (n = 10) and B) ADC-fMRI (n = 12), visualized using Circos^104^. The gray matter ROIs and white matter ROIs are indicated by dark and light bars. GM-GM, GM-WM and WM-WM edges are depicted with green, blue and red lines, respectively. The ROIs are divided according to the right (R) and left (L) hemisphere. The thickness of the lines represents the FC strength.

BOLD-fMRI displayed stronger negative correlations than ADC-fMRI. It is known that task-positive networks (such as the dorsal attention network -DAN-) negatively correlate with task-negative networks (such as the DMN)^44,45^. The main regions of the DAN include the intraparietal sulcus (the supramarginal gyrus -SMG- and postcentral gyrus -PoG- here), the frontal eye field (the MFG and precentral gyrus -PrG- here), the middle temporal region (the middle temporal gyrus -MTG- here), as well as white matter tracts such as the superior longitudinal fasciculus (SLF) and the splenium of the corpus callosum (sCC)^46–48^. We found that the MTG and the superior temporal gyrus (STG) negatively correlated with the ACgG for both BOLD-fMRI and ADC-fMRI, and that the MTG negatively correlates with the MSFG for ADC-fMRI.

At 7T, similar positive correlations were visible in ADC-fMRI and BOLD-fMRI (Supplementary Fig. 1). The negative correlations were greatly attenuated in ADC-fMRI, but appeared strongly in BOLD-fMRI, especially in the GM-GM. The negative correlations also appeared between the MTG and the MSFG for ADC-fMRI, and between the STG and the ACgG/MSFG for both ADC-fMRI and BOLD-fMRI.

### GM-WM connectivity

At 3T, in ADC-fMRI, the sCC (DAN) was negatively correlated with the ACgG and the MFC (DMN) (Supplementary Fig. 11), but positively correlated with the PCgG and the PCu (DMN) (Fig. 2, blue edges, and Supplementary Fig. 11). In BOLD-fMRI, the sCC was also positively correlated with the PCgG. Although non-significant, the correlations sCC-MFC and sCC-PCu of BOLD-fMRI had the same polarity. In ADC-fMRI, the SLF (DAN) negatively correlated with the FRP and the MSFG (DMN). Although non-significant, the correlations SLF-FRP and SLF-MSFG of BOLD-fMRI had the same polarity. Both ADC-fMRI and BOLD-fMRI showed anti-correlations between the SLF and the MFC. The SLF was also positively correlated with the PCu (DMN) in ADC-fMRI (as well as in BOLD-fMRI, although non-significant). Consistent with Ding et al. ^22^, the tapetum (TAP) was negatively correlated with the MSFG in BOLD-fMRI, while the anterior corona radiata (ACR) was positively correlated with the ACgG in both contrasts. It should be noted that the mean of significant GM-WM FC weights was significantly larger (p = 0.0336) with the downsampled BOLD-fMRI than ADC-fMRI (Fig. 3 A vs Supplementary Fig. 8A) At 7T, the genu of the corpus callosum (gCC) and the ACR were positively correlated with the ACgG in both contrasts (Supplementary Fig. 1). Finally, the cingulum (hippocampus) (CgH) was negatively correlated with the triangular part of inferior frontal gyrus (TrIFG) in BOLD-fMRI.

**Figure 3:**
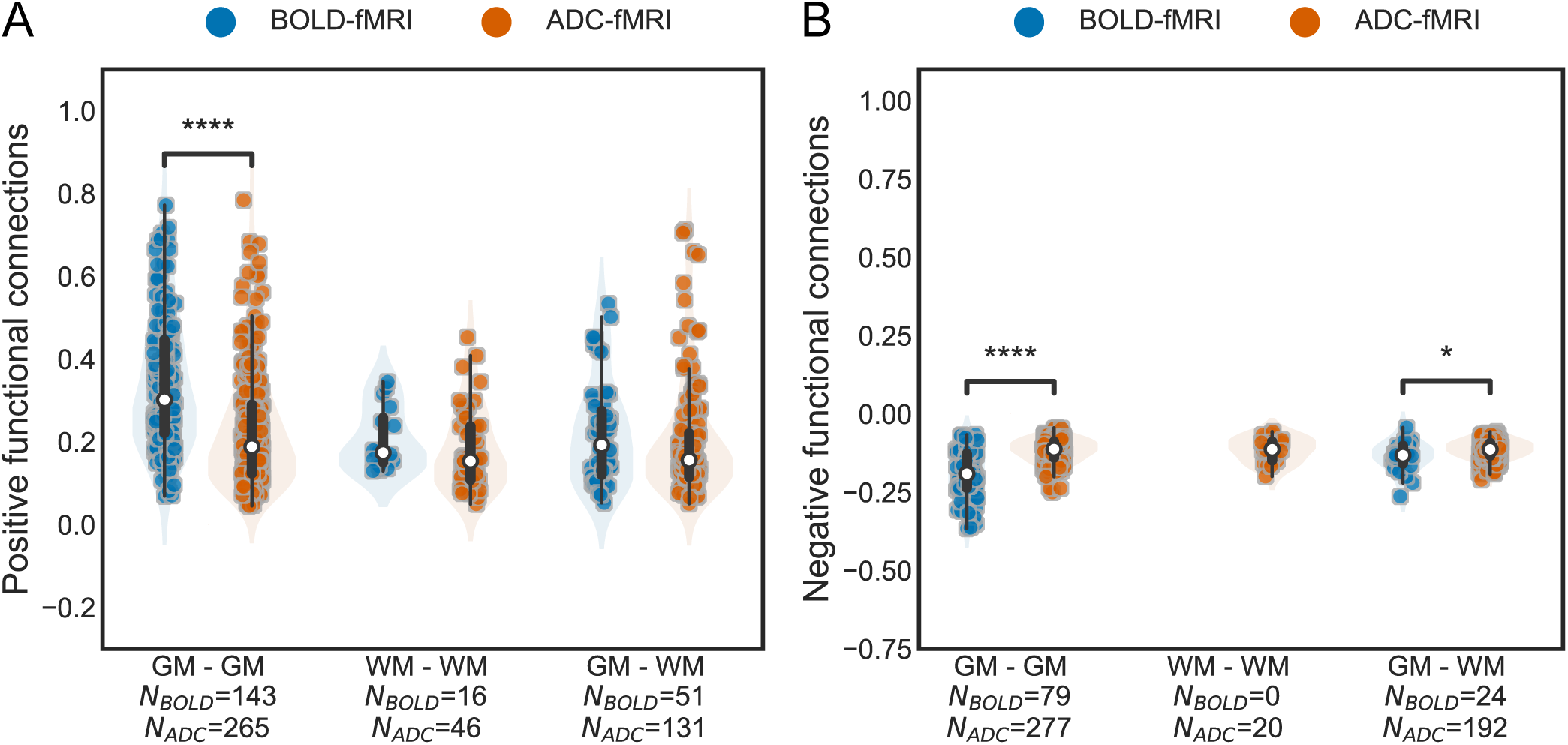
Significant positive and negative group-averaged functional connections. The group-averaged correlations are split by tissue type (GM-GM, WM-WM, and GM-WM connectivity). Each dot corresponds to a significant edge of the FC, averaged across subjects. A) Positive correlations, B) negative correlations. The number of significant edges per functional contrast is indicated on the x-axis label. No negative correlations were found for BOLD-fMRI WM-WM. A box-and-whisker plot is shown within each violin plot, with a white dot for the median. Two-sided Mann-Whitney-Wilcoxon tests with Bonferroni correction are performed. * 0.01 *<* p ≤ 0.05; ** 0.001 *<* p ≤ 0.01; *** 0.0001 *<* p ≤ 0.001; **** p ≤ 0.0001.

### WM-WM connectivity

At 3T, in both contrasts, the gCC was positively correlated with the ACR (Fig. 2, red edges, and Supplementary Fig. 11). In addition, the posterior thalamic radiation (PTR) was positively correlated with the SLF. The median FC strength decreased from 0.30 to 0.17 (43%) and 0.19 to 0.15 (21%) between GM-GM and WM-WM, for BOLD-fMRI and ADC-fMRI, respectively (Fig. 3).

At 7T, the same association between gCC and ACR was found (Supplementary Fig. 1). With ADC-fMRI, positive FC between the external capsule (EC) and the anterior/posterior limb of the internal capsule (ALIC/PLIC) and between the fornix (cres)/stria terminalis (FX/ST) and the sagittal stratum (SS) were also found. The median FC strength decreased from 0.38 to 0.20 (47%) and 0.28 to 0.24 (14%) between GM-GM and WM-WM, for BOLD-fMRI and ADC-fMRI, respectively (Supplementary Fig. 3).

### ADC vs BOLD connectivity strength

To further check the congruence between ADC-fMRI and BOLD-fMRI, we compared the connectivity strength of ADC-fMRI edges with the corresponding BOLD-fMRI edges, as proposed by Olszowy et al. ^25^, but this time extending the analysis to GM-WM and WM-WM edges (Fig. 4).

**Figure 4:**
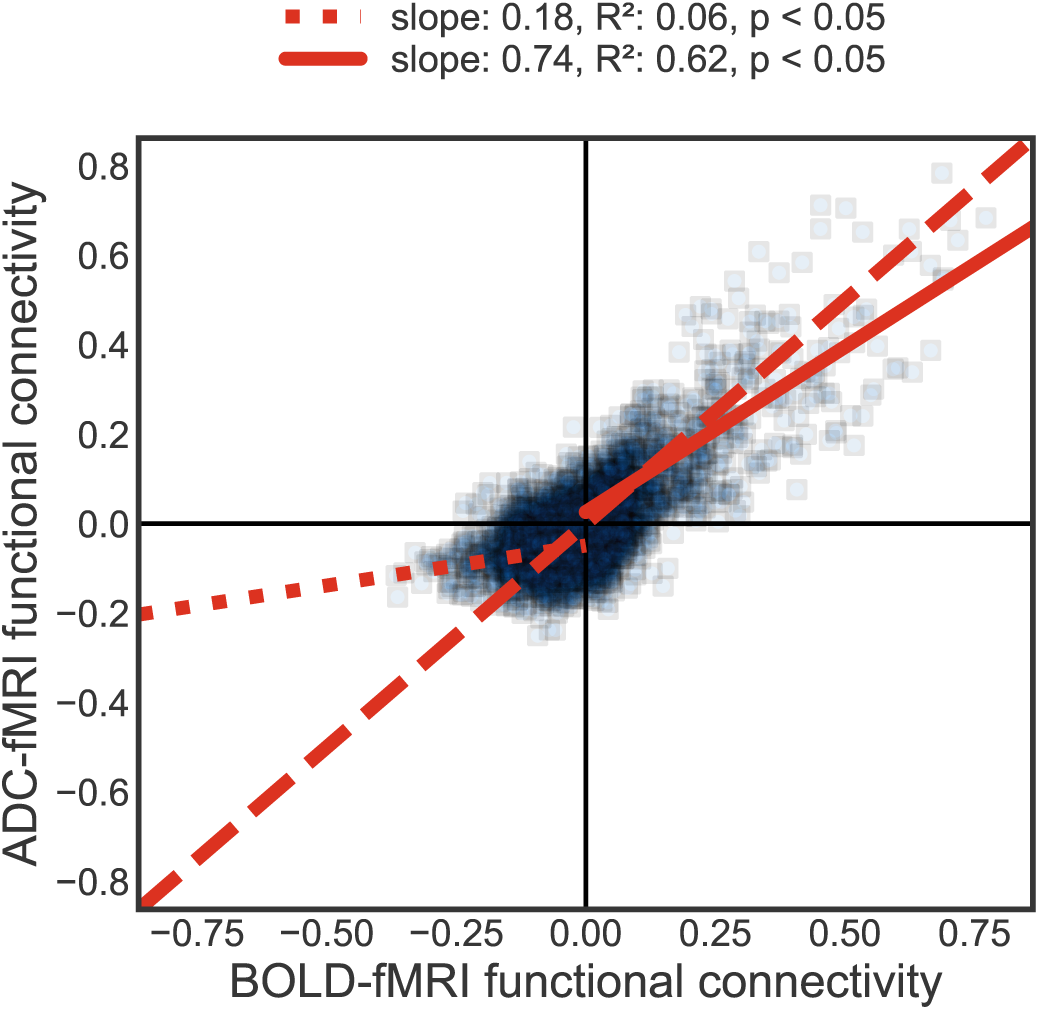
ADC-to-BOLD agreement of mean FC strength, between corresponding edges. The solid line fits the positive correlations, while the dotted line fits the negative correlations. The dashed line represents a perfect agreement (identity line). Functional connectivity strength is defined as Fisher-transformed Pearson’s correlation coefficient. There is excellent agreement between ADC-fMRI and BOLD-fMRI positive FC, while BOLD-fMRI negative FC is largely suppressed in ADC-fMRI.

For the positive functional connections, the linear fit between ADC-fMRI and BOLD-fMRI had a slope of 0.74 and a *R*^2^ of 0.62. For the negative functional connections, the fit had a slope of 0.18 but a *R*^2^ of 0.06. The comparison of connectivity strength between ADC-fMRI and BOLD-fMRI without GSR can be found in Supplementary Fig. 13. Without GSR, the excellent agreement between ADC-fMRI and BOLD-fMRI positive FC remained, while the disagreement in negative FC was not possible to assess anymore due to a near absence of negative edges.

The same analysis was conducted at 7T with similar results (Supplementary Fig. 2).

To gain more specific insight into the distribution of positive and negative FC in the GM-GM, WM-WM and GM-WM connectivity, we further segmented the data based on these specific domains (Fig. 3).

At 3T, positive FC was indeed significantly stronger (p *<* 0.0001) for BOLD-fMRI than for ADC-fMRI in the GM-GM, supporting the visual inspection (Fig. 3A). In WM-WM and in GM-WM, no differences were found in positive connections between the two contrasts. For the negative connections, they were significantly attenuated in ADC-fMRI in the GM-GM (p *<* 0.0001) and GM-WM (p = 0.0288) (Fig. 3B), but no anti-correlations were found for BOLD-fMRI in the WM-WM. Comparison of BOLD to ADC connectivity strength per domain and without GSR can be found in Supplementary Fig. 14. Without GSR, the results in positive GM-GM FC in Fig. 3A were preserved, while negative FC could not be evaluated as negative edges were almost non-existent.

At 7T, similar results are found, except that the WM-WM positive connections were significantly greater (p = 0.0225) for ADC-fMRI vs BOLD-fMRI (Supplementary Fig. 3A) and the negative edges were not significantly attenuated in the GM-WM anymore (Supplementary Fig. 3B).

At both field strengths, the number of significant positive and negative edges detected with ADC-fMRI was larger than with BOLD-fMRI.

### Graph Analysis

Following the analysis of individual FC edges, we then studied the networks formed by the significant positive GM-GM and WM-WM edges of the FC matrices (Fig. 1B). Weighted metrics (average node strength, average clustering and global efficiency) were computed and compared between the two contrasts.

At 3T, in the GM-GM, the average clustering and average node strength were similar between ADC-fMRI and BOLD-fMRI (Fig. 5A). They were however significantly larger (p = 0.0003 for average clustering, p = 0.0007 for average node strength) for ADC-fMRI than BOLD-fMRI for networks built on WM-WM connectivity (Fig. 5B). The global efficiency was significantly larger for ADC-fMRI in both the GM-GM (p = 0.0041) and WM-WM (p *<* 0.0001) connectivity. Graph analysis results without GSR can be found in Supplementary Fig. 15. They showed similar results, except that the global efficiency was not significantly different between BOLD-fMRI and ADC-fMRI in either connectivity domain.

**Figure 5:**
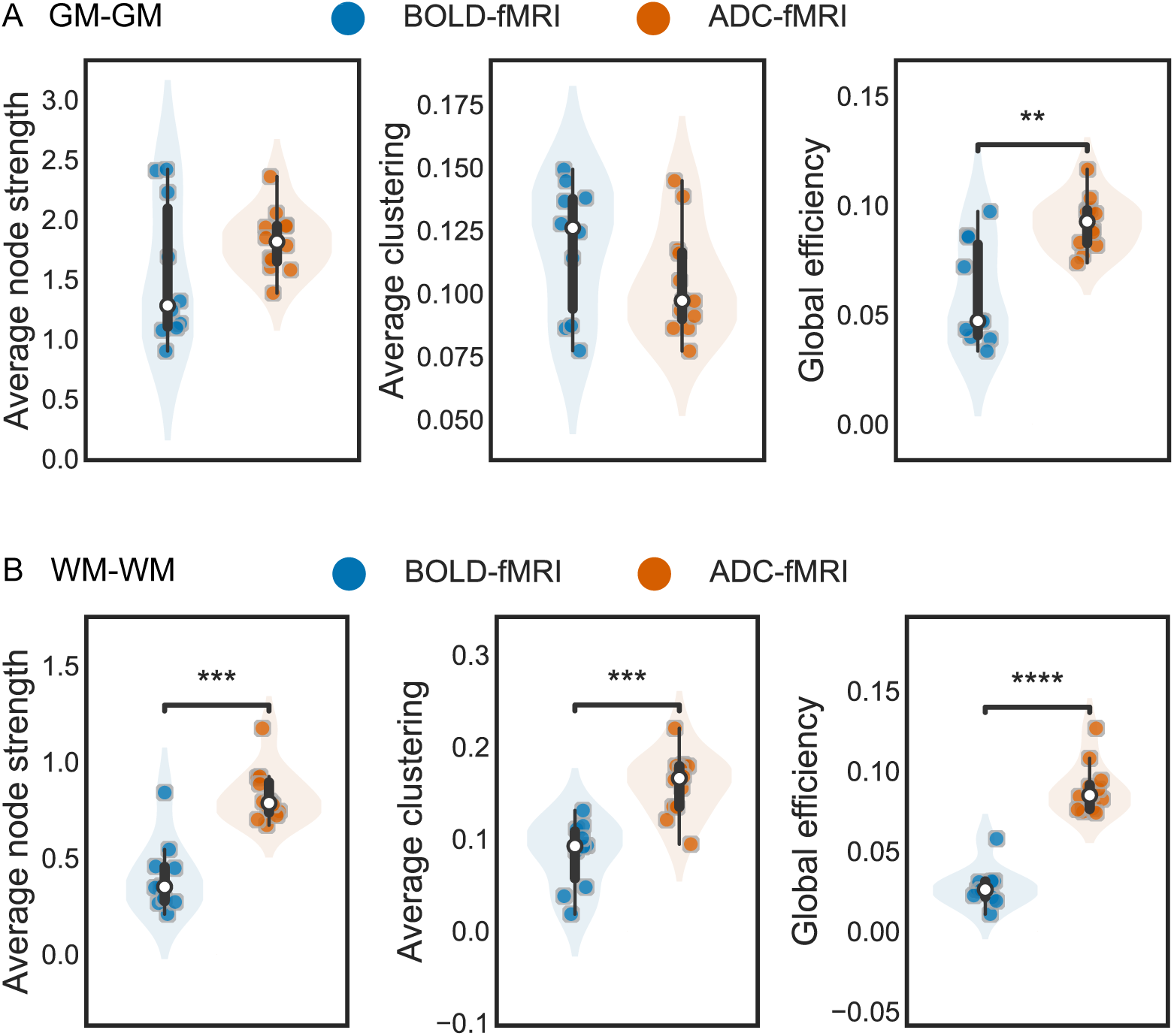
Subject-wise weighted graph metrics. Graph metrics of BOLD-fMRI (n = 10) and ADC-fMRI (n = 12), based on A) GM-GM connectivity, or B) WM-WM connectivity. A box-and-whisker plot is shown within each violin plot, with a white dot for the median. Two-sided Mann-Whitney-Wilcoxon tests with Bonferroni correction were performed. * 0.01 *<* p ≤ 0.05; ** 0.001 *<* p ≤ 0.01; *** 0.0001 *<* p ≤ 0.001; **** p ≤ 0.0001.

At 7T, the average node strength and global efficiency were similar in the GM-GM (Supplementary Fig. 4A), but became significantly larger (p = 0.0173 for average node strength, p = 0.0173 for global efficiency) in ADC-fMRI in the WM-WM (Supplementary Fig. 4B). The average clustering was significantly larger (p = 0.0376) in BOLD-fMRI in the GM-GM, but significantly larger (p = 0.0010) in ADC-fMRI in the WM-WM.

### Inter-subject similarity

Finally, we assessed the consistency of FC patterns across subjects by computing inter-subject similarity matrices (Pearson correlations between the significant edges, excluding self-connections, Fig. 1B) for both ADC-fMRI and BOLD-fMRI data, in the different connectivity domains (Fig. 6).

**Figure 6:**
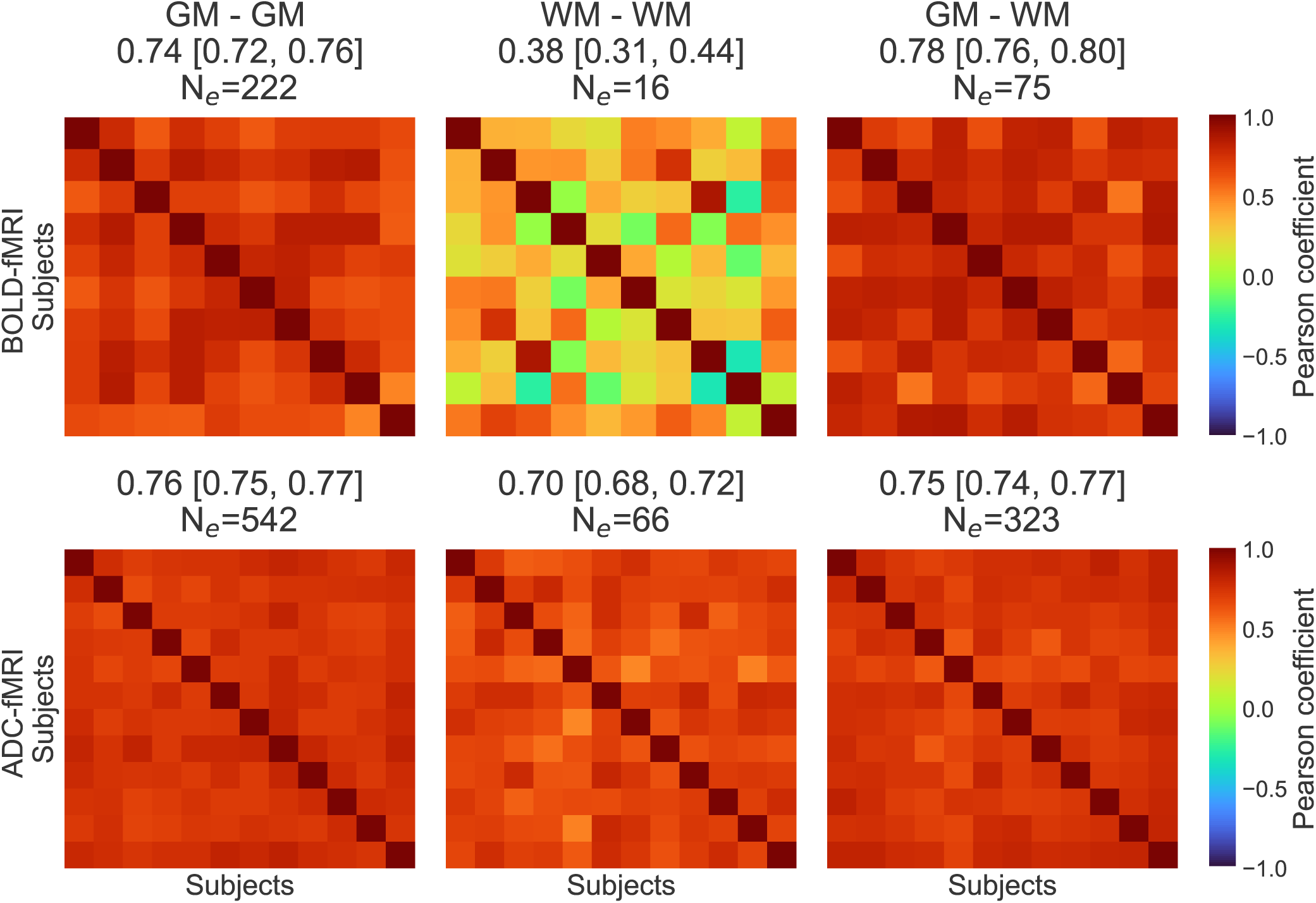
Inter-subject FC similarity of the significant edges. The inter-subject FC similarity is calculated as the pairwise Pearson correlation of the vectorized FC matrices. Each similarity matrix has a size N*_subjects_* x N*_subjects_*, with N*_BOLD_*_−_*_fMRI_* = 10 and N*_ADC_*_−_*_fMRI_* = 12. The mean Pearson’s correlation coefficient of the similarity matrix and the 95% CI are depicted in the subtitles. N*_e_* corresponds to the number of FC significant edges on which the Pearson’s correlation coefficients are calculated. The GM-GM, WM-WM and GM-WM connectivity were considered separately.

At 3T, the consistency across subjects (measured as pairwise Pearson correlation of the vectorized FC matrices) was larger for ADC-fMRI (mean 0.70, 95% CI [0.68, 0.72]) than for BOLD-fMRI (0.38 [0.31, 0.44]) in the WM-WM. Note that the number of significant edges N*_e_* used to calculate the pairwise correlation between two subjects was systematically larger for ADC-fMRI than BOLD-fMRI. A similar analysis was therefore performed on all edges (N*_e_* = 210 in both contrasts) to ensure that the significant difference in the WM-WM inter-subject similarity between ADC-fMRI and BOLD-fMRI was not biased by the small sample size (Supplementary Fig. 16). In this case, the ADC-fMRI inter-subject similarity was still larger (0.52 [0.50, 0.55]) than the BOLD-fMRI similarity (0.42 [0.38, 0.47]). We also reported the inter-subject similarity in WM-WM for the downsampled BOLD-fMRI (0.56, CI [0.51, 0.60]) (Supplementary Fig. 10A).

At 7T, the inter-subject similarity was larger for ADC-fMRI than for BOLD-fMRI in all the connectivity regions (Supplementary Fig. 5). Consistent with the 3T findings, the mean WM-WM inter-subject similarity was 0.67 [0.64, 0.70] for ADC-fMRI, but 0.24 [0.18, 0.31] for BOLD-fMRI. The inter-subject similarity in WM-WM for the downsampled BOLD-fMRI is 0.20 [0.13, 0.27] (Supplementary Fig. 10B).

## Discussion

Functional MRI studies extending beyond GM to map WM activity have been gaining substantial interest^11,18,19^. Although resting-state FC that includes both gray and white matter regions has been occasionally studied using BOLD-fMRI^22–24^, this work is the first to assess ADC-fMRI resting-state FC in gray and white matter, and compare it to its BOLD-fMRI counterpart. We found that the positive FC was extremely well preserved between ADC-fMRI and BOLD-fMRI, while negative FC was heavily attenuated in ADC-fMRI. We further reported a higher number of significant edges in ADC-fMRI vs BOLD, across connectivity regions (GM-GM, GM-WM, WM-WM). Weighted graph metrics revealed significantly larger average node strength, average clustering and global efficiency for ADC-fMRI than BOLD-fMRI in the WM-WM domain, while in the GM-GM domain, only global efficiency was higher with ADC vs BOLD. Finally, the inter-subject similarity was also larger in ADC-fMRI vs BOLD-fMRI in WM-WM, suggesting a higher reproducibility of ADC-fMRI FC in that domain.

ADC-fMRI and BOLD-fMRI positive FC mirrored each other, both with their mean FC strength (as previously noted in Olszowy et al. ^25^) and their weighted graph metrics (average node strength and clustering). Overall, positive FC was larger in BOLD-fMRI than ADC-fMRI, due to its higher CNR, particularly in GM. We further extended this analysis to individual edges and found that the DMN stood out, as expected during a resting state acquisition. Forming the anterior DMN, the ACgG, the MSFG and the FRP have high FC to each other^49^. Moreover, positive FC was also found between the ACgG and the MFC^50^. Together, these established connections provide additional assurance of the high correspondence of positive functional connections between ADC-fMRI and BOLD-fMRI positive FC, and confirm that ADC-fMRI is a reliable tool to decipher resting-state connectivity in the GM.

Conversely, negative correlations are more difficult to interpret. In this work, we found negative FC between the MTG and the MSFG/ACgG for BOLD-fMRI, as previously described in a similar analysis also performed with GSR^51^. In addition, we observed BOLD negative correlations between the sCC and the ACgG/MFC, between the TAP and the MSFG, as well as between the CgH and the TrIFG, as reported in a study not using GSR^22^. We also found these negative correlations in ADC-fMRI, indicating that both neurovascular and neuromorphological contrasts are sensitive to these negative correlations and that the latter may have a physiological significance. Commonly reported negative correlations are between antagonist resting-state networks such as the DMN and the DAN^44,45^.

However, in our analysis, almost all negative correlations disappeared when performed without GSR. Earlier studies showed that negative correlations could be mainly GSR-driven^52,53^, introducing artefactual negative correlations, while other studies showed GSR to improve FC specificity when used with care^51,54–56^, despite only partially removing “physiological connectivity”^57^. Another explanation could be that, in the same way the global signal fluctuations mask established neuronal associations concerning positive correlations, they could also obscure negative correlations in BOLD-fMRI FC^51^. More and more studies are reporting their results with and without GSR, finding negative correlations in both cases^56,58^. In this work, they were largely absent or attenuated in ADC-fMRI in spite of an analysis pipeline identical to BOLD-fMRI, including GSR. If GSR was inducing negative correlations, we would expect it to cause similar changes to both ADC-fMRI and BOLD-fMRI. In particular, in Ding et al. ^22^, the analysis was performed without GSR, which suggests that the mentioned anti-correlations are not necessarily artifactual. While the larger positive FC of BOLD could be explained by larger CNR, the preferential attenuation of anti-correlations in ADC-fMRI vs BOLD-fMRI is beyond what would be expected from CNR differences in the positive case. Nor can it be explained by the different temporal resolution of the two contrasts, as the results still hold when BOLD-fMRI is downsampled to the same sampling rate as ADC-fMRI. This suggests that while some BOLD-fMRI negative correlations may be related to brain activity, some may also be purely driven by vasculature or physiological noise. Adding to this, in the WM, where the vasculature is largely reduced, no significant anti-correlations were found for BOLD-fMRI. Bianciardi et al. ^45^ supported this hypothesis by reporting that blood volume changes may also induce negative correlations, especially in periventricular regions, as well as sulcal areas next to large pial veins. Anti-correlations can however also be caused by noise sources such as motion or physiological variations.

Initially thought to be driven by physiological noise, the BOLD-fMRI WM signal is increasingly recognized as consisting of hemodynamic changes reflecting neural activity in the WM or in neighboring GM regions, although with reduced sensitivity^11,18,19^. Thus, a concern with GM-WM and WM-WM connectivity is that it may be contaminated by GM signal via partial volume effect at the tissue boundary. Even though partial volume effect at the tissue interface is unavoidable, it will contribute in a similar way to BOLD-fMRI and ADC-fMRI, since their spatial resolution, the parcellation used, and the registration process are the same. In addition, GM ROIs are gyrified, so we would actually expect GM-GM to be more affected by partial volume with subcortical WM than GM-WM or WM-WM which is computed on JHU ROIs in the deep WM (mostly disjoint from cortical GM). In this work, we compared for the first time BOLD-fMRI and ADC-fMRI derived GM-WM and WM-WM connectivity.

Few studies have considered GM-WM connectivity. Ding et al. ^22^ reported positive and negative BOLD-fMRI GM-WM correlations at 3T. Remarkably, our study revealed similar positive and negative BOLD-fMRI correlations to Ding et al. ^22^, Wakana et al. ^59^, Karababa et al. ^60^ at 3T (but also at 7T), which strengthens the viability of BOLD-fMRI to study ROI-based GM-WM connectivity. Moreover, most of the BOLD-derived GM-WM connectivity patterns were also found using ADC-fMRI. This suggests that ADC-fMRI is also a suitable tool to detect positive and negative FC in the GM-WM edges.

Li et al. ^20^ derived BOLD HRFs for superficial, middle and deep WM. They found that the BOLD HRF corresponding to superficial WM was similar to the BOLD GM HRF, possibly allowing correlation analysis between WM and GM timecourses. However, they reported that middle and deep WM HRFs could display an additional delay to peak, possibly due to the different nature and distance of the blood vessels^20^. Taken together, the heterogeneity of the WM HRFs, if confirmed, could bring an additional challenge to deriving GM-WM functional connectivity. We hypothesize that BOLD-derived WM-GM FC analysis will be mostly sensitive to the superficial WM.

Similarly to GM-WM, WM-WM connectivity has been scarcely studied^24^. In BOLD-fMRI, the magnitude is larger than in ADC-fMRI for both positive and negative FC, whereas in WM, the magnitude is smaller for both positive and negative FC. A possible explanation is that some of the BOLD negative correlations are purely vascular in origin (spurious signals). In the GM-GM connectivity, where the vasculature is dense, BOLD-fMRI has higher positive FC (possibly related to higher CNR) and higher negative FC (some of which may be spurious vascular signals). In the WM-WM, the vasculature is sparse, resulting in reduced low frequency oscillations, blood flow and blood volume^1,11^, and therefore reduced BOLD CNR. The positive FC in WM-WM (following brain activity) is still detected, albeit with lower amplitudes, but the negative FC in WM-WM is no longer detectable with BOLD-fMRI, possibly as a consequence of both the reduced CNR and the reduced sensitivity of this tissue to spurious vascular signals. As the SNR difference between GM and WM within contrasts is reasonably comparable, and the SNR is systematically larger in BOLD-fMRI than in diffusion-weighted time series (Supplementary Table 1), we do not expect it to be the cause of the observed difference. On the other hand, ADC functional contrast is assumed to be driven by neuromorphological coupling (or glia-morphology coupling) and thus has the potential to sustain similar CNR levels in WM-WM vs GM-GM FC.

Further looking into the graph metrics analysis, we found larger average clustering coefficients and average node strength in ADC-fMRI than in BOLD-fMRI in the WM-WM connectivity, suggesting the two functional contrasts capture different connectivity aspects in the WM. Global efficiency was larger in ADC-fMRI than in BOLD-fMRI in both WM-WM and GM-GM at 3T, but particularly in the WM-WM. Differences in graph metrics may be driven by ADC-fMRI robustness to vascular signal, and therefore possibly to physiological noise in the GM-GM, and by decreased BOLD-fMRI sensitivity in the WM-WM. Taken together, the ADC-fMRI network metrics reflected both better integration and segregation in the WM than BOLD-fMRI.

Because GSR can impact FC values by shifting them towards more negative values, it can ultimately impact network metric calculations. Indeed, the networks on which the metrics were calculated ignored negative FC values. This selection of positive FC edges may affect the BOLD-fMRI and ADC-fMRI graph metrics differently due to different distributions of positive and negative correlations. Therefore, we compared graph metrics derived from ADC-fMRI vs BOLD-fMRI FC also without GSR, but found similar results, as also observed in Li et al. ^24^. This suggests that the graph metric differences between the two functional contrasts are not driven by GSR-induced biases.

Remarkably, our data also showed that ADC-fMRI has a higher between-subject similarity than BOLD-fMRI in the WM-WM connectivity, although SE BOLD has higher inter-subject similarity than GE BOLD^61^. This suggests that ADC-fMRI is more consistent across subjects, possibly due to the suppression of the vascular signal and attenuation of physiological noise thanks to the “self-normalization” inherent to the ADC calculation from two consecutive diffusion-weighted images. It cancels out not only T_2_-weighting but also fluctuations slower than the TR, due to respiratory and cardiac motion, for example. In terms of thermal noise, BOLD-fMRI SNR is systematically larger than the SNR from the diffusion-weighted time series, in both GM and WM (Supplementary Table 1). Therefore, low BOLD SNR in the WM cannot explain the observed lower WM-WM inter-subject similarity for BOLD-fMRI than for ADC-fMRI. Furthermore, for GM-GM connectivity, the FC matrices and between-subject similarity are comparable between BOLD-fMRI and ADC-fMRI. In a comparable context of SNR differences between GM and WM within contrast, the between-subject similarity is therefore higher for ADC-fMRI than for BOLD-fMRI.

We also investigated the potential effect of BOLD-fMRI higher temporal resolution. Because the WM-WM inter-subject similarity is still significantly smaller than ADC-fMRI WM-WM at 3T (Supplementary Fig. 10A) and at 7T (Supplementary Fig. 10B), we concluded that the temporal resolution is not the cause of this difference. Alternatively, the lower BOLD-fMRI inter-subject similarity could reflect the fact that BOLD-fMRI is more sensitive to inter-subject differences and is able to detect individual fingerprinting^62^. The latter is however more unlikely, as the BOLD-fMRI similarity coefficients are high in edges other than WM-WM.

Taken together, ADC-fMRI revealed consistent patterns of WM-WM connectivity, making it a useful tool for investigating WM functional connectivity. Despite the limitation of acquiring only one diffusion encoding direction, thus optimizing the sensitivity to axonal swelling perpendicular to this direction and limiting the sensitivity to axons with other directions (see Discussion), the WM-WM results are encouraging. They should be further improved by using isotropic diffusion encoding, to sensitize the ADC contrast to axon bundles independent of their orientation.

As mentioned in the Introduction, the underlying mechanism of the ADC-fMRI contrast is debated. While its advocates support the consensus that ADC fluctuations are driven by activity-induced neuromorphological changes, its critics claim that this contrast is either contaminated by the hemodynamic response or driven by morphological changes induced by vascular dilation^63,64^. In this work, we have shown that ADC-fMRI does not mimic BOLD, but is indeed a separate contrast. Notably, negative FC patterns were distinct between ADC-fMRI and BOLD-fMRI, negative correlations being highly attenuated in ADC-fMRI. In addition, the WM connectivity captured by both contrasts was significantly different, both in terms of graph metrics, but also in terms of consistency across subjects. While ADC-fMRI WM-WM connectivity was consistent across subjects, BOLD-fMRI WM-WM connectivity varied much more. Thanks to the mitigation techniques implemented in this study, ADC-fMRI is much more robust to vascular contributions. The higher number of significant edges found in ADC-fMRI across connectivity regions (GM-GM, GM-WM, WM-WM) could also stem from robustness against vascular signals. At 3T, the higher number of subjects for ADC-fMRI (n = 12) than BOLD-fMRI (n = 10), probably also increased the number of significant edges found in ADC-fMRI. However at 7T, despite the same sample size (n = 10), ADC-fMRI still detected more significant edges than BOLD-fMRI in all connectivity regions.

Our results were therefore in line with previous studies that aimed to untangle neuromorphological from vascular effects, using targeted inhibition of key physiological processes following brain activity. Disrupting the hemodynamic response, aquaporin-4 blocker^36^ and nitroprusside^37^ were used in mice and rats, respectively. In both cases, the BOLD response was notably affected while the ADC was not. Regarding neuromorphological effects, cell swelling induced via hypotonic solution or ouabain was shown to decrease ADC at the tissue level^65^, albeit outside of physiological ranges. Conversely, Abe et al. ^42^ showed that the ADC increased under the effect of the neural swelling blocker furosemide during anesthesia while the local field potentials (LFPs) remained unaffected. Our results provided additional support for the non-vascular origins of ADC changes during brain activity. Preclinical studies suggested these changes could be driven by fluctuations in cell shapes and sizes, as ADC is so sensitive to the microstructure layout. Other mechanisms cannot however be entirely ruled out.

By paralleling the analysis at 3T and 7T, we wanted to verify the influence of the field strength on each of the two functional contrasts. The magnitude of the BOLD signal is known to scale with field strength^66,67^, whereas we hypothesized that the magnitude of the ADC response should be independent of field strength, if stemming from neuromorphological coupling. However, field strength dependencies are complex and affect each functional contrast in multiple ways. For example, the SNR and the ratio of physiological to thermal noise increase with field strength ^67^. In addition, due to the shortening of blood T_2_ with increasing field strength, the BOLD signal contains less direct intravascular component, and therefore has increased spatial specificity. In contrast, ADC-fMRI timecourses are less sensitive to vascular contributions but ADC FC analysis could benefit from higher SNR at higher field. However, mitigating vascular contributions to the ADC timecourse may be more difficult at 7T due to increased magnetic susceptibility^25^. In the context of rs-fMRI, our results were similar at 3T and 7T, but the dependence of ADC-fMRI on field strength may become apparent in task-fMRI.

The small sample size is one of the main limitations of this study. It diminished the statistical power of the analyses, highlighting the need for future studies with larger sample sizes. In addition, to optimize spatial and temporal resolution, only a section of the brain was scanned (a partial coverage which was further reduced when coregistering subjects to a common space due to slight variations in slab positioning), which limited the extent of functional connectivity that could be examined across the brain. Furthermore, the use of atlas-based ROIs rather than participant-based segmentation for GM and WM could introduce additional uncertainty at the boundary between GM and WM and add to the partial volume effect. However, this is affecting BOLD-fMRI and ADC-fMRI similarly, as they share the same spatial resolution. In addition, the use of two b-values to calculate one ADC-fMRI time point resulted in a lower temporal resolution (2 s) for ADC-fMRI, possibly leading to the aliasing of faster physiological processes, whether related to brain activity or to other signals such as respiration and heart signal^68^. Also, DW-TRSE-EPI used in this study was only acquired with one diffusion encoding direction. This possibly yields an inhomogeneous sensitivity to neuromorphological changes across the brain, depending on the angle between the main fibers at each location and the diffusion encoding direction, with higher sensitivity for fibers perpendicular to that direction and lower for fibers parallel to that direction^43,69^. Note that, as GM is much less organized than WM, we expect this limitation to only affect the WM activity detection. Lastly, even though the use of SE-EPI rather than GE EPI sequence enabled a fairer comparison with the DW-TRSE sequence, SE BOLD exhibits less sensitivity than GE BOLD, possibly missing some of the resting-state BOLD characteristics described in the literature. However, SE BOLD has been previously shown to give very similar results to GE for rs-fMRI analyses^61^.

To overcome the partial brain coverage, additional acceleration in diffusion MRI acquisition needs to be introduced to acquire a large number of slices within a short TR, possibly using novel undersampling strategies^70^. This would enable a more comprehensive comparison of GM-WM correlations as presented in Ding et al. ^22^, as well as WM-WM. In addition, the temporal resolution could be increased by using a sliding window to calculate an ADC at each TR, although this is expected to introduce temporal auto-correlations and may not yield more informative ADC timecourses. Another option would be to keep analyzing the diffusion-weighted signal timecourse (from a single b-value, instead of the ADC), bearing in mind that there will be BOLD/T_2_ contribution to the DW signal in the GM, but considering that contribution to be minimal in WM. This would double the temporal resolution and enable the observation of faster processes. Cardiac and respiratory signals could also be recorded to directly track the physiological contribution to the BOLD-fMRI and ADC-fMRI signals. Nonetheless, it is most valuable to develop a functional contrast that is suitable for both GM and WM tissues. Furthermore, in our experience with task fMRI, the ADC timecourse superseded the DW signal timecourse in detecting WM activity^69^. The value of ADC vs DW signal timecourses for WM resting-state connectivity may however be studied further in future work. In future studies, isotropic diffusion encoding will be used to sensitize ADC-fMRI to fibers no matter their orientation^69^. While it could be interesting to investigate anisotropic diffusion effects linked to action potential propagation (“DTI fMRI”), earlier works that have looked at directional dependency of the ADC response reported maximal response perpendicularly to fibers and weak to absent change along fibers^43,71,72^. Simulations in realistic WM substrates have also shown much reduced sensitivity along axons vs transversally^69^. Furthermore, the low temporal resolution of a DTI fMRI experiment is for now dissuasive for resting-state fMRI. However, ADC-fMRI FC could be used in combination with structural tractography as a way to track brain activity along white matter fibers, up to the cortical areas. Lastly, since the ADC response is linked to microstructural changes that occur on a much faster timescale than the BOLD hemodynamic response, the advent of faster MRI techniques in combination with ADC-fMRI may allow the observation of such faster processes.

This study indicates that ADC-fMRI is a promising tool for resting-state analysis. It retained comparable positive correlations to BOLD-fMRI in the GM. Most interestingly, ADC-fMRI showed clearly distinct functional contrast from neurovascular coupling, notably via largely reduced anti-correlations in the GM. Furthermore, larger average clustering coefficients, average node strength and global efficiency in the WM-WM connectivity as compared to BOLD-fMRI, combined with higher inter-subject similarity in the WM-WM, suggest that ADC-fMRI has a high sensitivity and robustness to WM functional connectivity. In the future, it holds great potential for elucidating WM functional connectivity as an alternative to BOLD-fMRI. Future work will use an isotropic diffusion encoding to sensitize the ADC-fMRI in all directions, thus preventing bias towards fibers perpendicular to the diffusion encoding direction.

## Methods

### Data acquisition

The study was approved by the Ethics Committee of the Canton of Vaud (CER-Vaud) and the dataset was previously reported on in Olszowy et al. ^25^. Twenty-two subjects (7 males, age 25 ± 5) were scanned after giving their informed written consent. All ethical regulations relevant to human research participants were followed. Siemens Prisma 3T and Siemens Magnetom 7T scanners (Siemens, Erlangen, Germany), equipped with 80 mT m^−1^ gradients, were used to acquire the data. A 64-channel and a 32-channel receiver head coils were used at 3T and 7T, respectively. Subjects were scanned at 3T and 7T in order to examine any potential field strength dependency of the ADC-fMRI functional contrast, as compared to BOLD-fMRI.

For anatomical reference and atlas registration, a whole brain 1-mm isotropic T_1_-weighted image was acquired using a MP-RAGE sequence (matrix size: 224 x 240 x 256).

We used a DW-TRSE-EPI sequence for ADC-fMRI and a SE-EPI sequence for BOLD-fMRI. As briefly mentioned in the Introduction, the TRSE sequence was chosen to minimize cross-terms with background field gradients. It should be noted that the cross-term suppression assumes constant background field gradients^73^. In fMRI, the background field gradients due to blood susceptibility changes are not constant, but can be considered to be slowly varying on the timescale of the diffusion encoding. In this context, the effect of cross-terms in the diffusion weighting is thus not suppressed, but nonetheless attenuated by the use of bipolar gradient TRSE design. SE BOLD was employed instead of the conventional gradient echo (GE) BOLD to allow for a more comparable design to DW-TRSE-EPI. Indeed, any residual BOLD contribution to the ADC-fMRI timecourses would be more similar in origin to the SE BOLD contrast. Note that, despite reduced sensitivity, the use of SE instead of GE for BOLD largely cancels out contributions from extravascular water around macrovasculature, increasing BOLD spatial specificity to capillaries^61,74–76^. However, one cannot assume that the macrovascular effect has completely disappeared. During the long echo train, only the echo at TE is perfectly refocused. This introduces some additional T^∗^ weighting in the MR signals, which results in a larger relative signal change (ΔS/S) during functional activation than expected with a pure T_2_-weighted SE BOLD^77,78^. Despite its reduced sensitivity, SE BOLD was shown to result in comparable FC maps to GE BOLD at 3T^61^, and to yield typical resting-state networks at 7T^79^, thus demonstrating the validity of using SE BOLD contrast for rs-fMRI.

For rs-fMRI runs, participants were instructed to fixate on a cross in the center of a screen and to relax. Two rs-fMRI scanning protocols were used: (1) 2D multi-slice SE-EPI (T_2_-BOLD contrast) and (2) 2D multi-slice DW-TRSE-EPI with alternating pairs of b-values 200 and 1000 s mm^−2^. Acquisition parameters common to both protocols were: TE: 73 ms (3T) or 65 ms (7T), TR: 1 s, matrix size: 116 x 116, 2-mm in-plane resolution, 16 x 2-mm slices, in-plane GRAPPA acceleration factor 2^80^, partial Fourier 3/4, simultaneous multi-slice (multiband) factor 2^81,82^, 600 volumes for a scan time of 10 min. The pairs of alternating b-values in the DW-TRSE-EPI sequence were combined to give a resulting ADC-fMRI time series with a sampling period of 2 s. The BOLD-fMRI results presented in this paper were calculated from time series with a TR of 1 s but this higher temporal resolution may affect the comparison between BOLD-fMRI and ADC-fMRI. Therefore, the potential effect of the different temporal resolution on the results was investigated by deriving the results on BOLD-fMRI time series downsampled to a sampling period of 2 s, keeping every other time point (see Supplementary Fig. 6-10), and comparing them with the results presented in this paper. The multi-band accelerated EPI sequences (http://www.cmrr.umn.edu/multiband/) were provided by the Center for Magnetic Resonance Research (CMRR) at the University of Minnesota^83^. The main acquisition parameters are summarized in Supplementary Table 2.

The relatively short TR used to achieve reasonably high temporal resolution only enabled partial brain coverage (32-mm thick slab). The imaging slab (Supplementary Fig. 12) was placed to encompass part of the DMN, including the prefrontal cortex and the posterior cingulate cortex^4,25^. A few EPI volumes (N = 4) with opposite phase encoding direction were acquired for susceptibility distortion correction.

### Data processing

The data were processed as described in Olszowy et al. ^25^. (1) Slices at the extremities of the slab were excluded from the analysis due to inflow contributions^76^. (2) The volume outliers were automatically detected (signal dropouts) and replaced with linearly interpolated signals. For a time point to be considered as an outlier, the linearly detrended average brain signal should deviate by more than 1% (3T) or 3% (7T) from the median of the detrended signal time series. This step aims to discard volume outliers that are due to drastic events (mainly motion). Thresholds were experimentally set to detect these extreme outliers at each field strength. Despite different expected fluctuation amplitudes between BOLD-fMRI and ADC-fMRI, a whole brain fluctuation larger than 1% or 3% is a very large global effect in both cases. If the proportion of outlier time points exceeded 20%, the entire run was excluded from the analysis. For ADC-fMRI, this processing was carried out separately on b_1_ and b_2_ time series. The fraction of outliers did not exceed 5% in most of the cases. (3) MP-PCA denoising with a sliding window kernel of 5 x 5 x 5 voxels was applied on the data to boost the tSNR^84–86^. For ADC-fMRI, it was conducted separately on the two b-values. The residuals were checked for Gaussianity, confirming that the denoising did not alter data distribution properties. (4) Gibbs ringing correction was applied^87^. (5) Susceptibility distortions^88^ were corrected with FSL *topup*, using the settings from Alfaro-Almagro et al. ^89^. (6) Motion artifacts were corrected with SPM^90^. For ADC-fMRI, motion correction was performed separately on b_1_ and b_2_ time series, followed by alignment of the two series using ANTs rigid transformation^91^. (7) For ADC-fMRI, ADC time series were obtained following Eq. 1. No spatial smoothing was used. (8) Independent component analysis (ICA) in FSL MELODIC^92^ with high-pass temporal filtering (f *>* 0.01 Hz) and 40 independent components was used to manually identify and remove structural noise (such as motion or physiological noise) components^85,93^. (9) Average BOLD-fMRI and ADC-fMRI timecourses were calculated in each ROI in native space using atlas-defined ROIs, described below (Fig. 1A).

### Atlas registration

Brain extraction was performed using ANTs on the T_1_-weighted volume before registration to the Montreal Neurological Institute (MNI) 1 mm isotropic template using symmetric normalization with ANTs. Using rigid transformation with ANTs, the average volume of each functional run was registered to the T_1_-weighted anatomical volume. The quality of the partial brain data registration (EPI onto T1w) was visually inspected in three planes (axial, sagittal, coronal) and considered to be good. For ADC-fMRI, the average functional volume was calculated on the b_1_ = 200 s mm^−2^ volumes only. The brain segmentation atlas was transformed to the subject’s space using the inverse of the two previous transformations (MNI → T_1_ → functional volume). The atlas used was a combination of the 1.5 mm isotropic Neuromorphometrics (NMM) gray matter atlas (*Neuromorphometrics, Inc.*, 136 ROIs) and the John Hopkins University (JHU) 1 mm isotropic white matter atlas (https://identifiers.org/neurovault.collection:264, 48 ROIs).

The use of a morphometric rather than a functional atlas is motivated by the fact that current functional atlases are derived from BOLD-fMRI, and we aimed to select a parcellation that was independent of the either functional contrast under investigation. Furthermore, comparing structural and functional data can be informative, as there is an expected relationship between the two. Due to partial brain coverage, only ROIs present in all the subjects, in both contrasts (BOLD and ADC), and containing at least ten voxels were retained (Supplementary Tables 3 and 4).

### Functional Connectivity

BOLD-fMRI and ADC-fMRI FC matrices were obtained by calculating Pearson’s partial correlation between each pair of atlas ROI timecourses, using the average timecourse of the whole brain (the global signal) as a covariate, followed by Fisher’s transform (Fig. 1A). Although it is hypothesized that the ADC signal is only sensitive to localized brain activity and therefore should not be affected by the global signal regression (GSR), we decided to apply the same processing to both BOLD-fMRI and ADC-fMRI, to allow a fair comparison between the two contrasts. Furthermore, residual physiological noise in ADC could be further mitigated by GSR. FC matrices were finally averaged across subjects for each contrast. Since GSR may also remove relevant functional signal and alter the distribution of correlation coefficients^94^, we duplicated the BOLD-fMRI and ADC-fMRI analyses without GSR to test the robustness of the results. Results without GSR are shown in the Supplementary Fig. 13, 14, and 15.

### Graph analysis

Only significant positive correlations in individual FC matrices were considered (Fig. 1B), determined with two-sided one-sample Wilcoxon tests with multiple comparisons correction (FDR Benjamini-Hochberg^95^). Weighted metrics for testing the network strength (average node strength), segregation (average clustering) and integration (global efficiency) were calculated for each subject. The detailed formula of the weighted metrics can be found in Rubinov and Sporns ^96^.

The node strength corresponds to the sum of the connection weights for a given node (the weighted equivalent of the node degree), averaged across all nodes to yield average node strength for the network^96^. The clustering coefficient of a node is a measure of inter-connectivity (local functional connectivity) between its neighboring nodes^97–99^. The clustering coefficient of a network (averaged across nodes) indicates the level of clustered connectivity around individual nodes, thus it is a measure of network segregation^96,97^.

The global efficiency is the average of the inverse shortest path length between all pairs of nodes, where path lengths are defined inversely to FC. Shorter paths between nodes indicate a high possibility of information flow across the brain, therefore a higher efficiency corresponds to enhanced potential for integration^96^.

### Statistics and reproducibility

The regression lines between FC values derived from each contrast were measured for positive and negative group-mean edge weights separately.

Significant positive and negative FC weights within each contrast were determined with two-sided one-sample Wilcoxon tests with multiple comparisons correction (FDR Benjamini-Hochberg^95^). These significant connections were then split according to their connectivity domains (GM-GM, WM-WM, GM-WM). To assess differences between ADC-fMRI and BOLD-fMRI, we compared significant FC edges in each domain using two-sided Mann-Whitney-Wilcoxon tests, with Bonferroni correction for multiple comparisons.

We also compared the ADC-fMRI and BOLD-fMRI graph metrics in the GM-GM and in the WM-WM domains using two-sided Mann-Whitney-Wilcoxon tests, with Bonferroni correction for multiple comparisons.

Finally, the pairwise inter-subject similarity within each contrast was derived by calculating the Pearson correlation coefficients between the significant GM-GM, WM-WM, or GM-WM edges, excluding self-connections (Fig. 1B).

In all statistical tests, *α* = 0.05.

The results were successfully replicated between field strengths (3T vs 7T), but the dataset used does not include test-retest scans.

## Data availability

The raw data are available in the openneuro repository https://openneuro.org/datasets/ds003676^100^. Data supporting Fig. 3-6, and Supplementary Fig. 11 and 16 are available in the figshare repository https://doi.org/10.6084/m9.figshare.28435325.v2101.

## Code availability

The Matlab code used to process the data is available on https://github.com/Mic-map/DfMRI ^102^. The python code used to run the analyses is available at https://github.com/Mic-map/ADC rsfMRI^103^.

## Acknowledgements

The authors thank Thomas Bolton and Jasmine Nguyen-Duc for insightful discussions. They acknowledge the CIBM Center for Biomedical Imaging for providing expertise and resources to conduct this study.

This work was supported by the Swiss National Science Foundation under a Spark award CRSK-2 190882 and by the Swiss Secretariat for Research and Innovation (SERI) under an ERC Starting Grant award ‘FIREPATH’ MB22.00032.

## Author contributions

Conceptualization: I.J., I.d.R.; Methodology: I.J., I.d.R., A.S.; Validation: I.d.R.; Formal analysis: I.d.R.; Investigation: W.O.; Data curation: W.O., I.d.R.; Writing-original draft: I.d.R.; Writing - Review & Editing: I.d.R., I.J., W.O., A.S.; Visualization: I.d.R., A.S.; Supervision: I.J.; Funding acquisition: I.J.

## Competing Interests

The authors declare no competing interests.

## Supplementary Materials

**Supplementary Figure 1:**
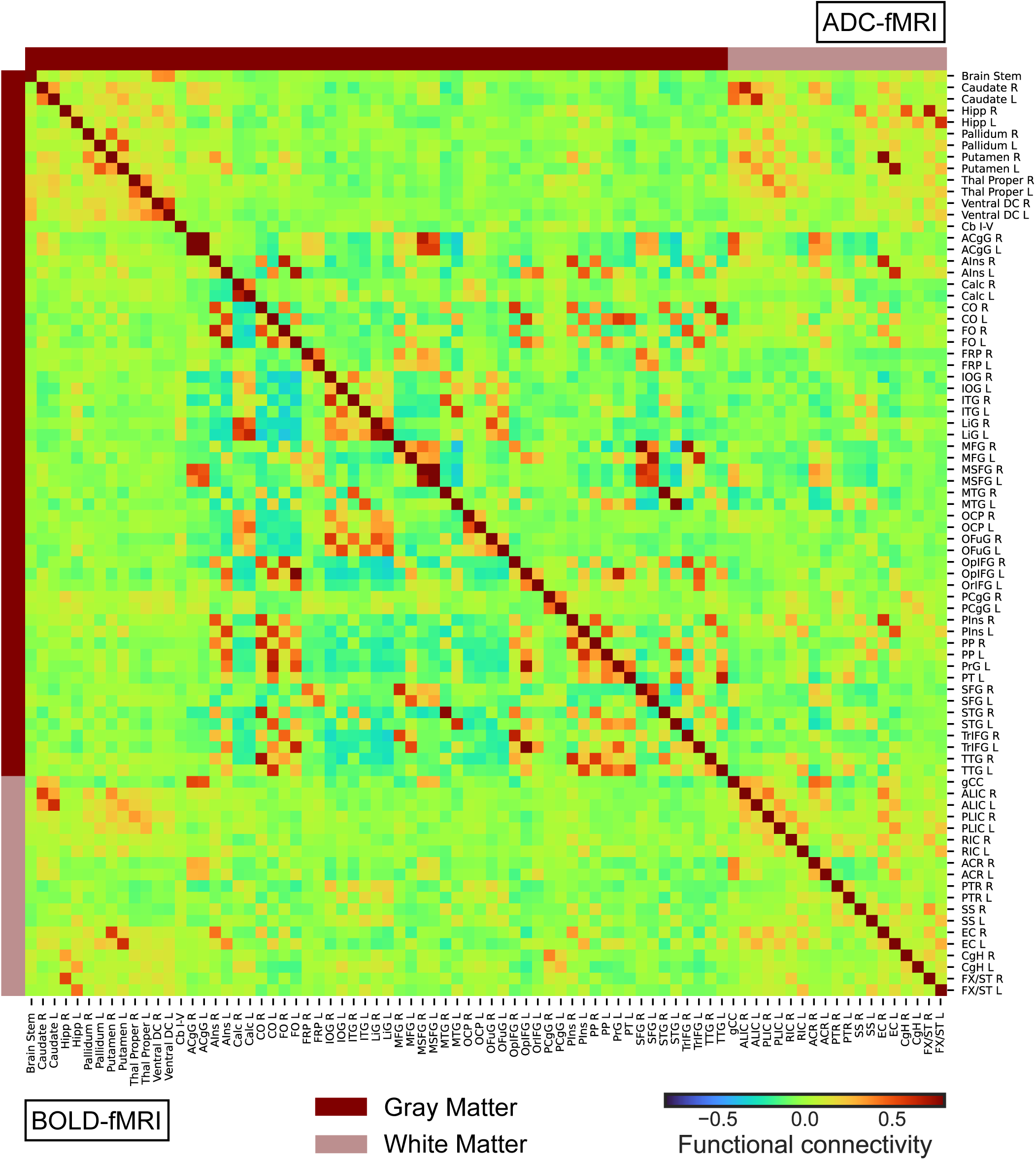
**7T: Mean Functional Connectivity**, for BOLD-fMRI (n = 10, lower triangle) and ADC-fMRI (n = 10, upper triangle). The gray matter ROIs and white matter ROIs are indicated by dark and light bars. Functional connectivity, defined as Fisher-transformed Pearson’s correlation coefficient, is indicated by the color bar.

**Supplementary Figure 2:**
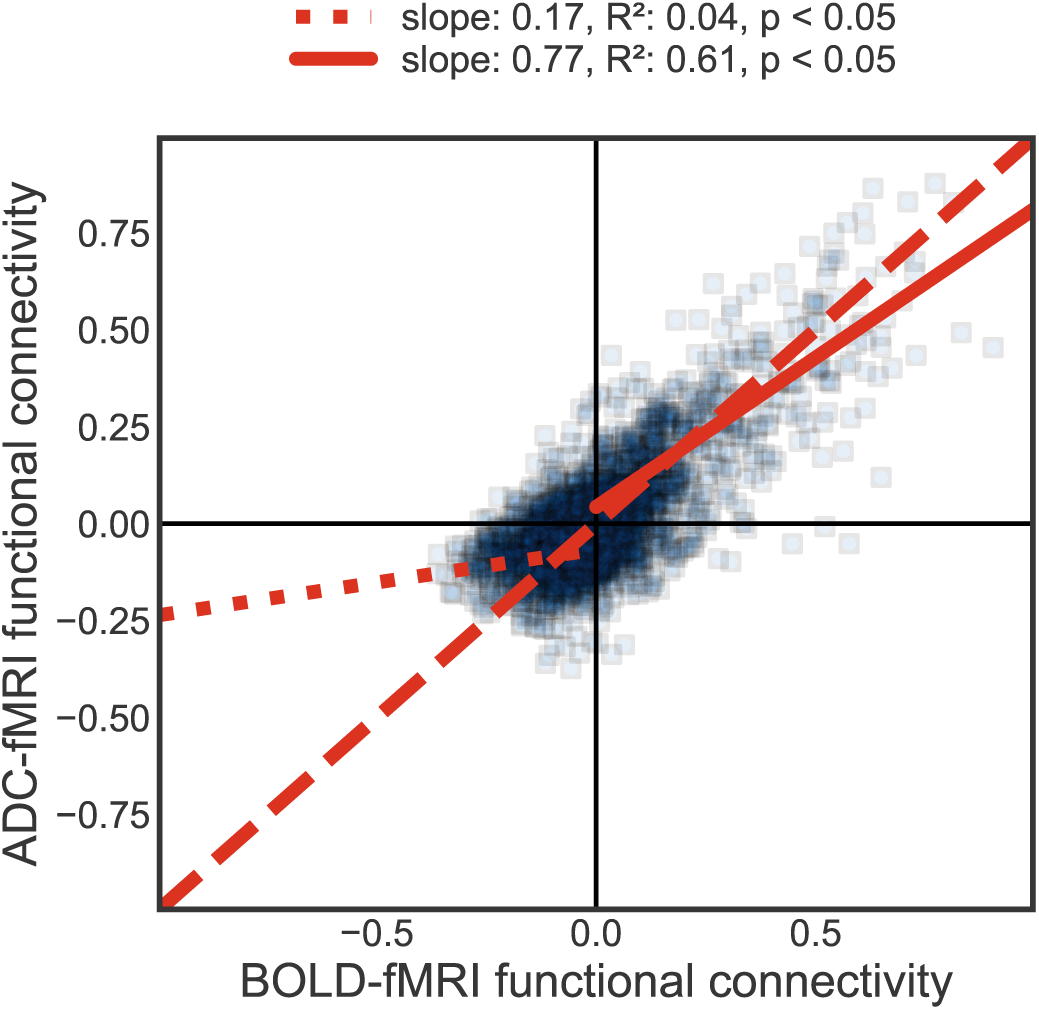
**7T: ADC-to-BOLD agreement of mean FC strength**, between corresponding edges. The solid line fits the positive correlations, while the dotted line fits the negative correlations. The dashed line represents a perfect agreement (identity line). Functional connectivity strength is defined as Fisher-transformed Pearson’s correlation coefficient. There is excellent agreement between ADC-fMRI and BOLD-fMRI positive FC, while BOLD-fMRI negative FC is largely suppressed in ADC-fMRI.

**Supplementary Figure 3:**
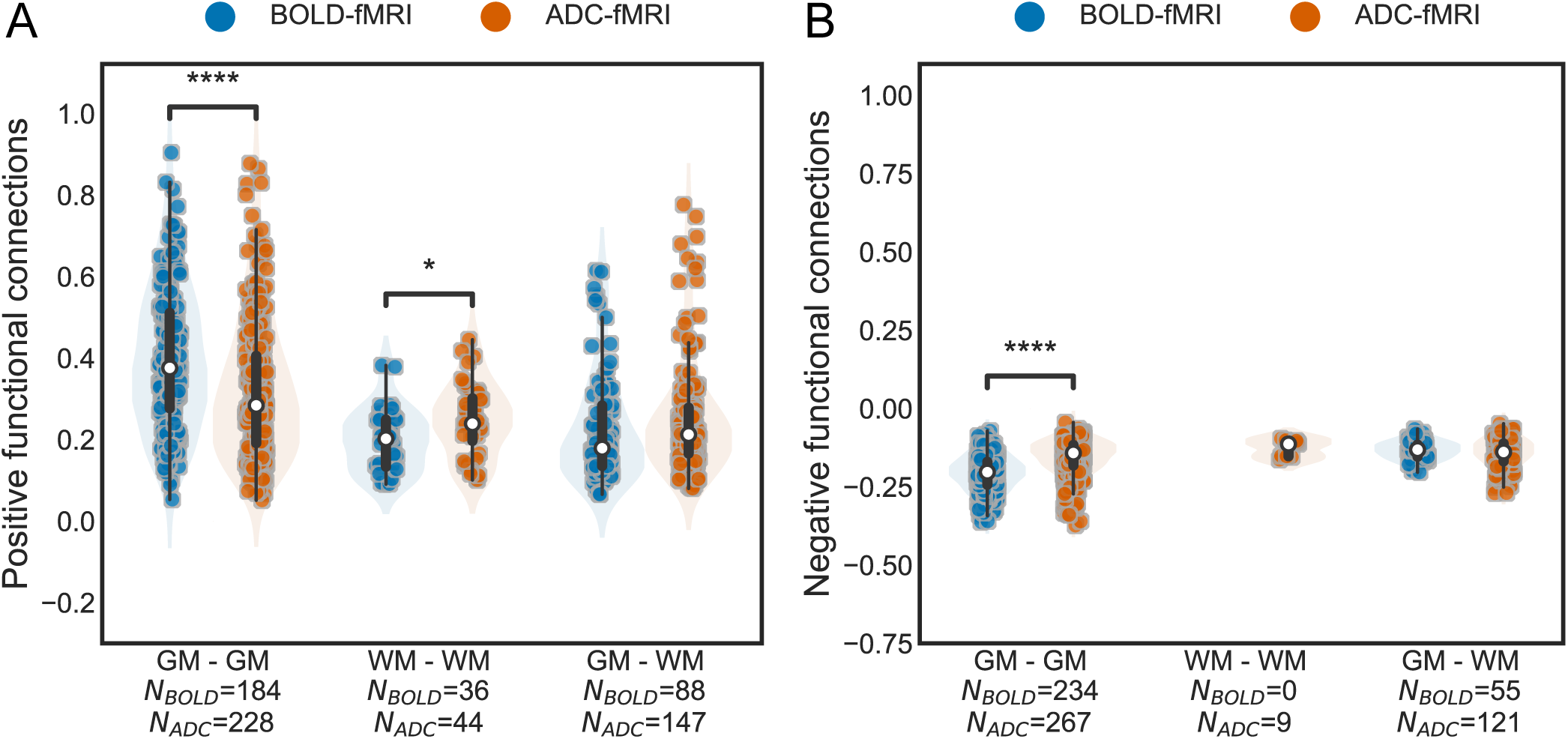
7T: Significant positive and negative group-averaged functional connections. The group-averaged correlations are split by tissue type (GM-GM, WM-WM, and GM-WM connectivity). Each dot corresponds to a significant edge of the FC, averaged across subjects. A) Positive correlations, B) negative correlations. The number of significant edges per functional contrast is indicated on the x-axis label. No negative correlations were found for BOLD-fMRI WM-WM. A box-and-whisker plot is shown within each violin plot, with a white dot for the median. Two-sided Mann-Whitney-Wilcoxon tests with Bonferroni correction are performed. * 0.01 *<* p ≤ 0.05; ** 0.001 *<* p ≤ 0.01; *** 0.0001 *<* p ≤ 0.001; **** p ≤ 0.0001.

**Supplementary Figure 4:**
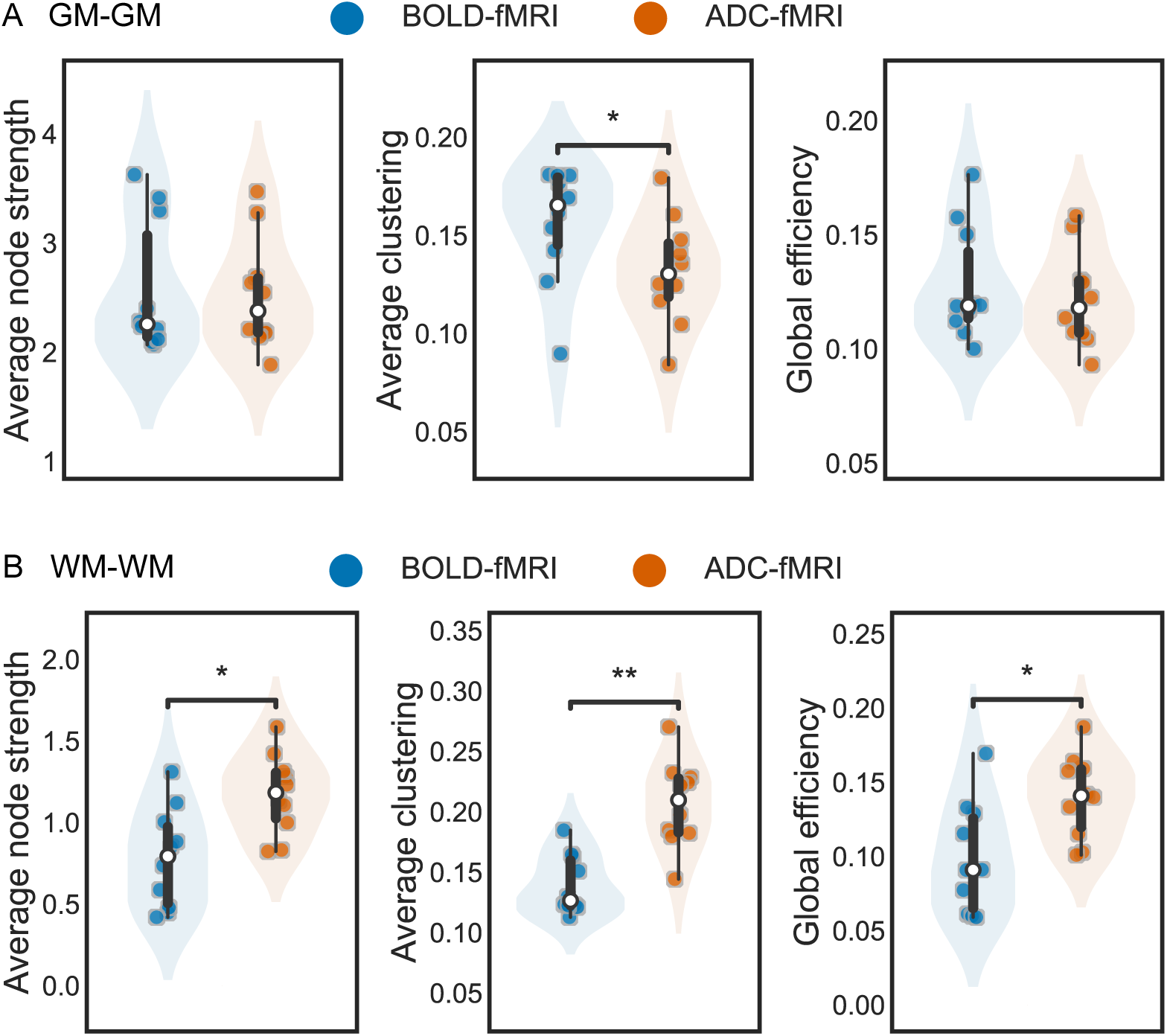
7T: Subject-wise weighted graph metrics. Graph metrics of BOLD-fMRI (n = 10) and ADC-fMRI (n = 10), based on A) GM-GM connectivity, or B) WM-WM connectivity. A box-and-whisker plot is shown within each violin plot, with a white dot for the median. Two-sided Mann-Whitney-Wilcoxon tests with Bonferroni correction were performed. * 0.01 *<* p ≤ 0.05; ** 0.001 *<* p ≤ 0.01; *** 0.0001 *<* p ≤ 0.001; **** p ≤ 0.0001.

**Supplementary Figure 5:**
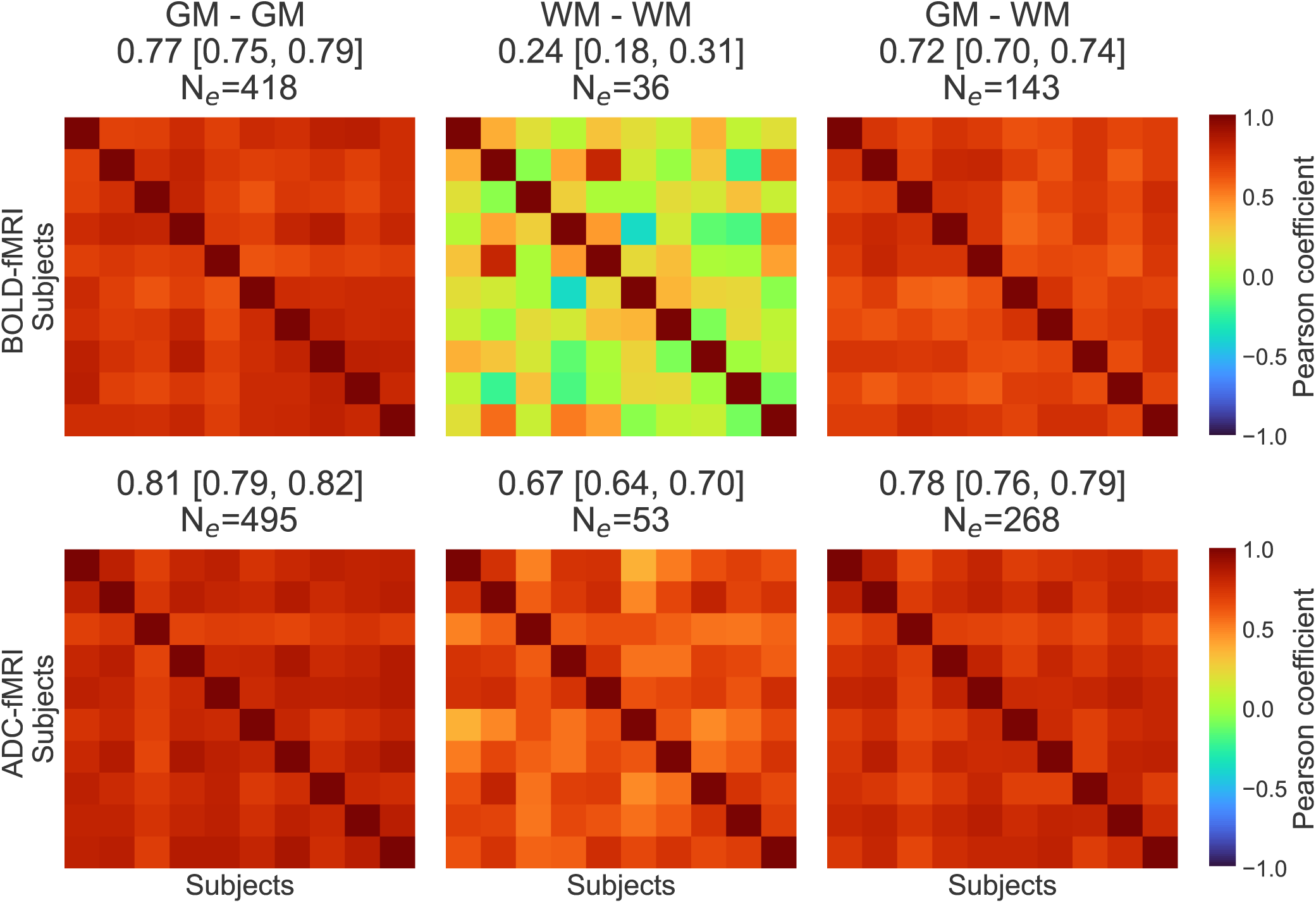
**7T: Inter-subject FC similarity** of the significant edges for BOLD-fMRI and ADC-fMRI, calculated as the pairwise Pearson correlation of the vectorized FC matrices. Each similarity matrix has a size N*_subjects_* x N*_subjects_*, with N*_BOLD_*_−_*_fMRI_* = 10 and N*_ADC_*_−_*_fMRI_* = 10. The mean Pearson’s correlation coefficient of the similarity matrix and the 95% CI are depicted in the subtitles. N*_e_* corresponds to the number of FC significant edges on which the Pearson’s correlation coefficients are calculated. The GM-GM, WM-WM and GM-WM connectivity were considered separately.

**Supplementary Figure 6:**
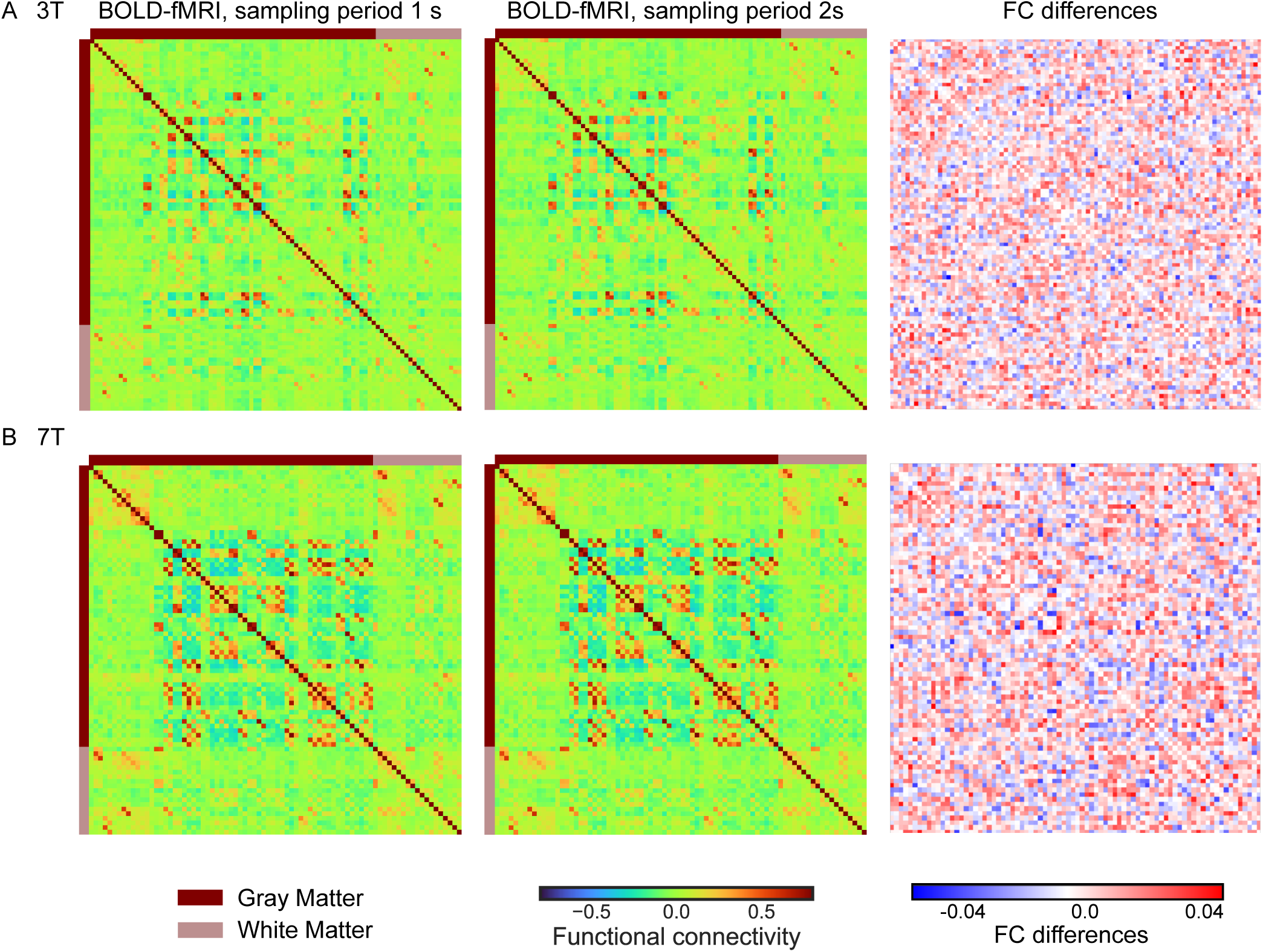
Downsampled BOLD: FC differences between BOLD with a sampling period of 1 s vs 2 s. A) 3T, B) 7T.

**Supplementary Figure 7:**
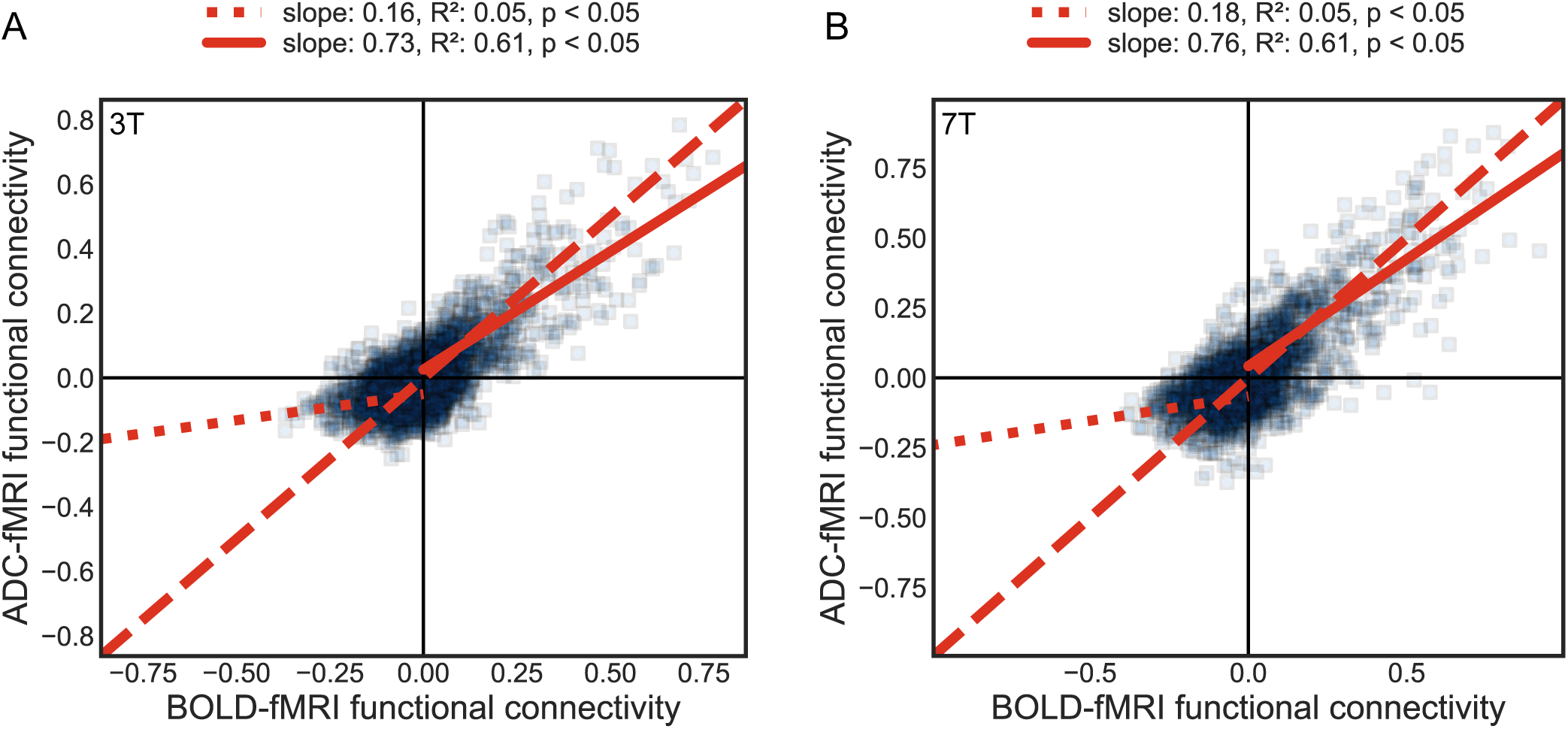
**Downsampled BOLD: ADC-to-BOLD agreement of mean FC strength**, between corresponding edges, for BOLD-fMRI and ADC-fMRI with matched sampling period. A) 3T, B) 7T. The solid line fits the positive correlations, while the dotted line fits the negative correlations. The dashed line represents a perfect agreement (identity line). Functional connectivity strength is defined as Fisher-transformed Pearson’s correlation coefficient.

**Supplementary Figure 8:**
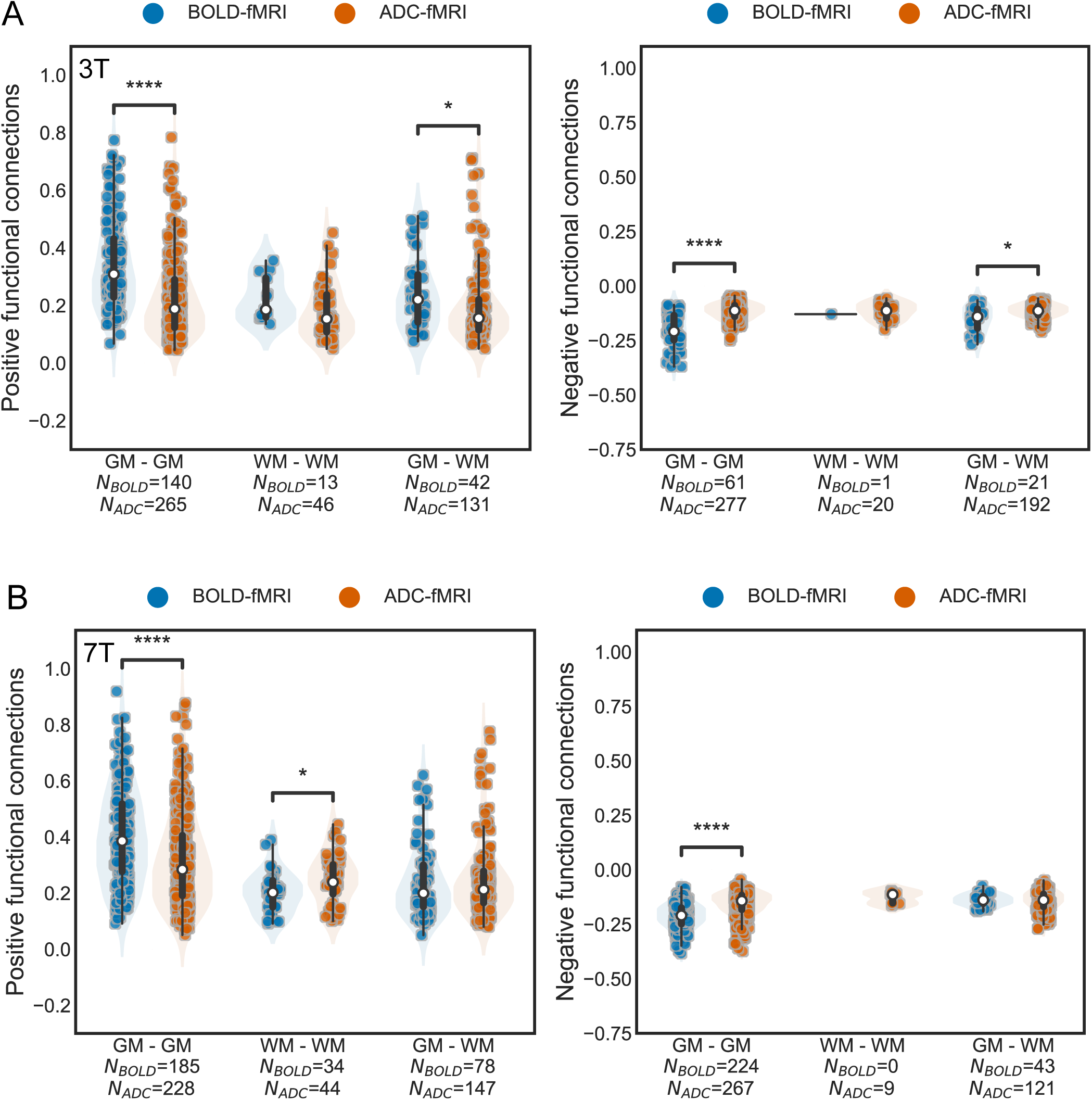
**Downsampled BOLD: Significant positive and negative group-averaged functional connections**, for BOLD-fMRI and ADC-fMRI with matched sampling period. BOLD time series were downsampled from a sampling period of 1 s to 2 s. The group-averaged correlations are split by tissue type (GM-GM, WM-WM, and GM-WM connectivity). Each dot corresponds to a significant edge of the FC, averaged across subjects. A) 3T, B) 7T. The number of significant edges per functional contrast is indicated on the x-axis label. A box-and-whisker plot is shown within each violin plot, with a white dot for the median. Two-sided Mann-Whitney-Wilcoxon tests with Bonferroni correction are performed. * 0.01 *<* p ≤ 0.05; ** 0.001 *<* p ≤ 0.01; *** 0.0001 *<* p ≤ 0.001; **** p ≤ 0.0001.

**Supplementary Figure 9:**
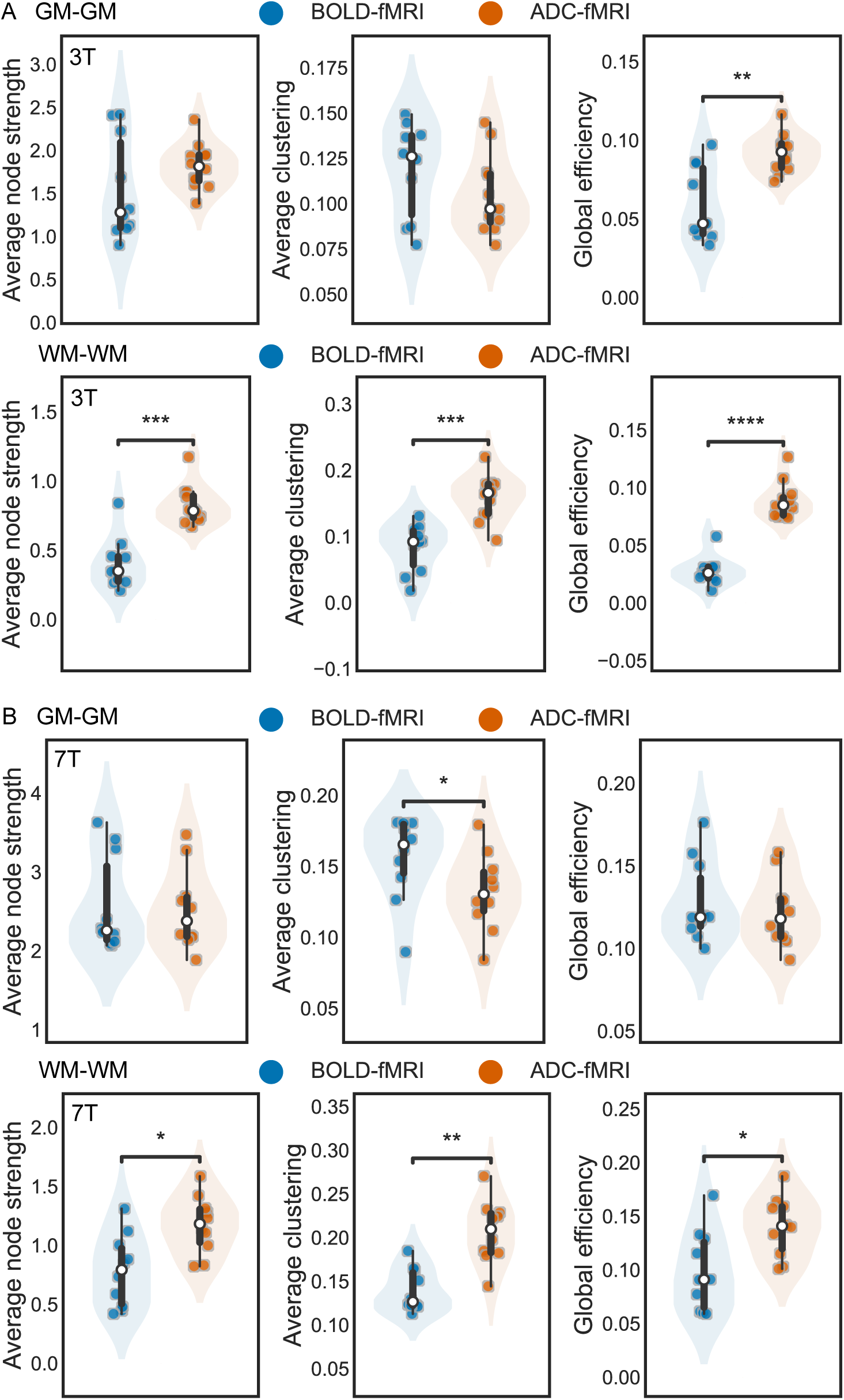
**Downsampled BOLD: Subject-wise weighted graph metrics**, for BOLD-fMRI and ADC-fMRI with matched sampling period. A) 3T, B) 7T. A box-and-whisker plot is shown within each violin plot, with a white dot for the median. Two-sided Mann-Whitney-Wilcoxon tests with Bonferroni correction were performed. * 0.01 *<* p ≤ 0.05; ** 0.001 *<* p ≤ 0.01; *** 0.0001 *<* p ≤ 0.001; **** p ≤ 0.0001.

**Supplementary Figure 10:**
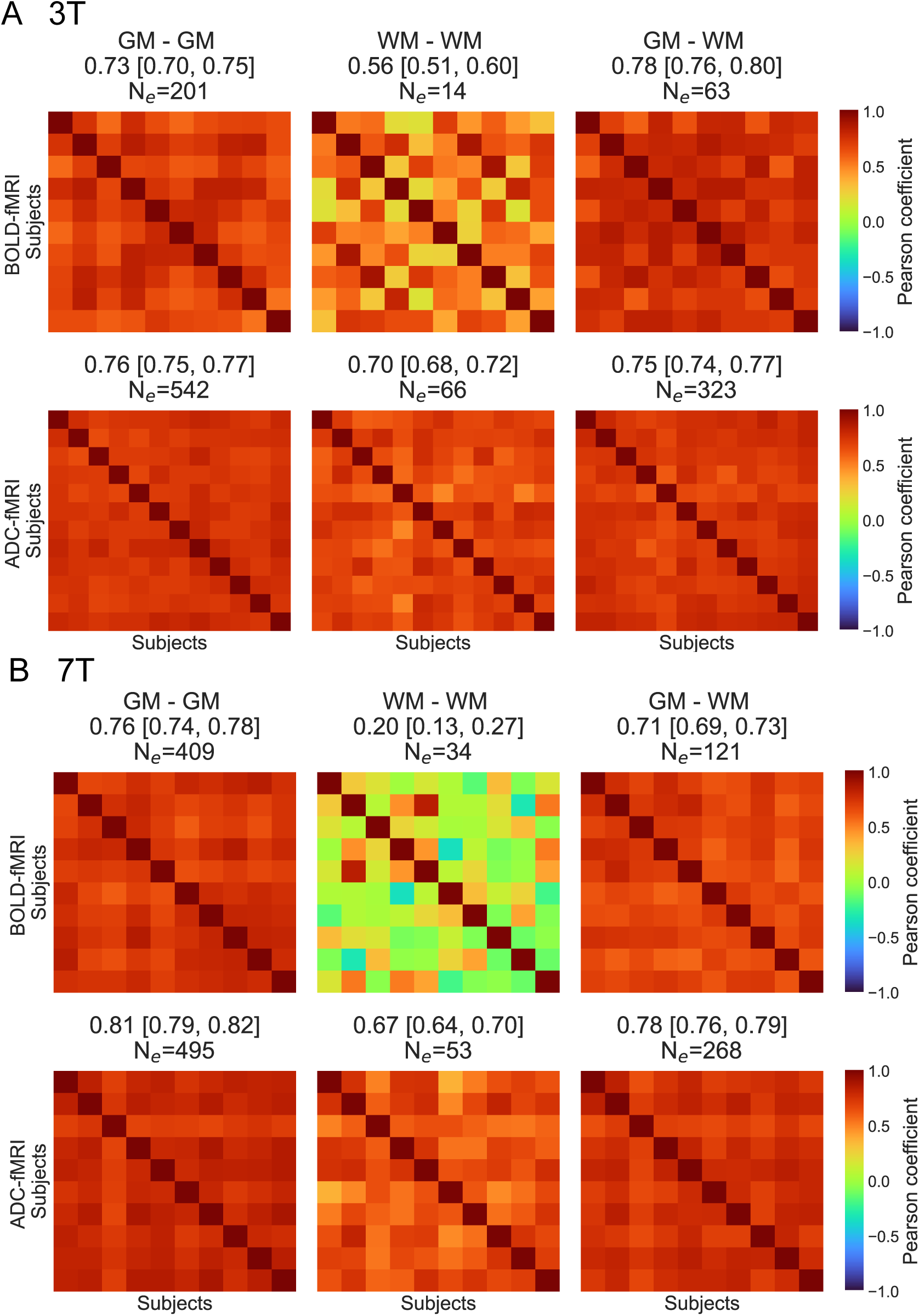
**Downsampled BOLD: Inter-subject FC similarity** of the significant edges for BOLD-fMRI and ADC-fMRI with matched sampling period, calculated as the pairwise Pearson correlation of the vectorized FC matrices. Each similarity matrix has a size N*_subjects_* x N*_subjects_*. The mean Pearson’s correlation coefficient of the similarity matrix and the 95% CI are depicted in the subtitles. N*_e_* corresponds to the number of FC edges on which the Pearson’s correlation coefficients are calculated.

**Supplementary Figure 11:**
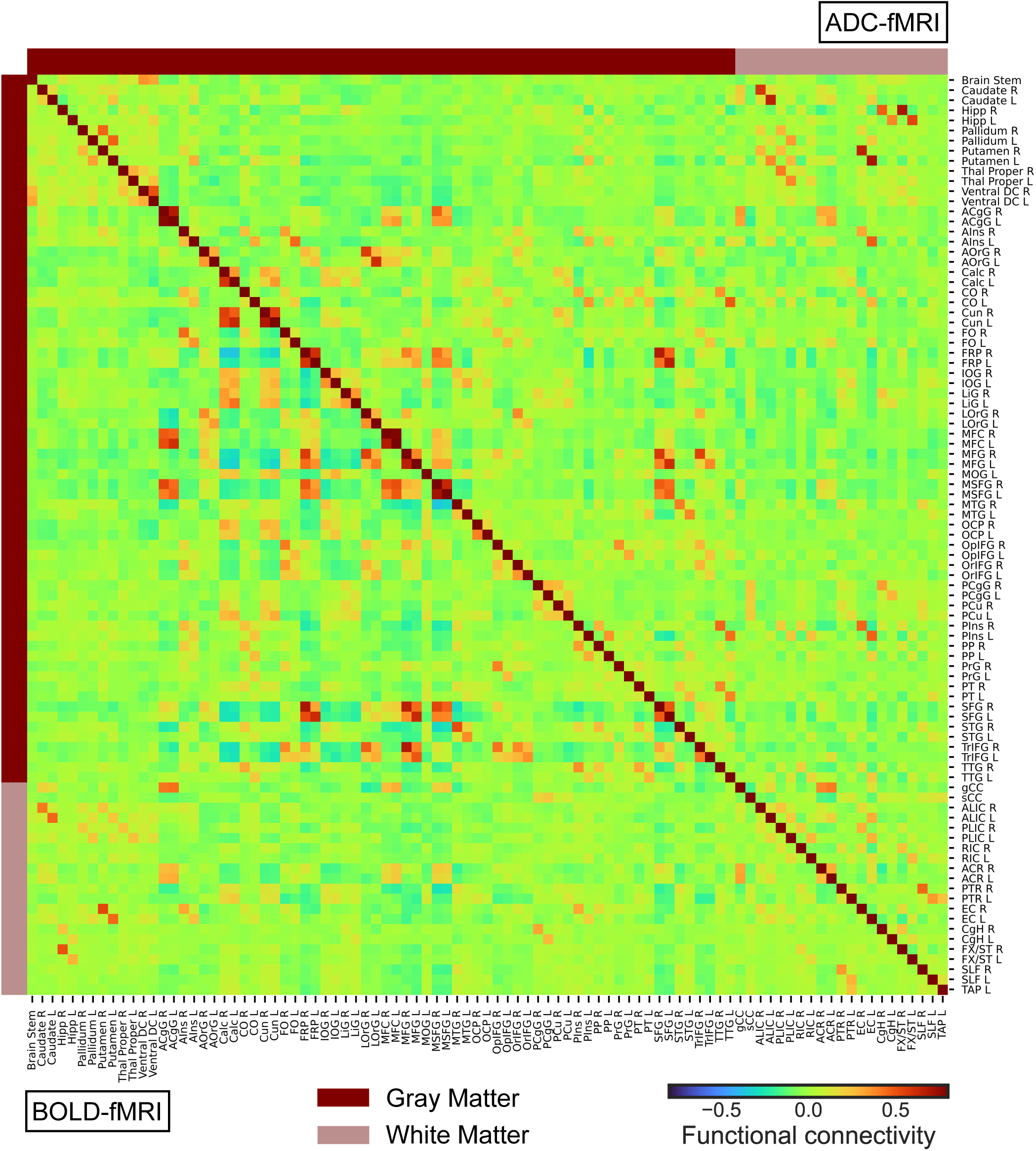
**3T: Mean Functional Connectivity**, for BOLD-fMRI (n = 10, lower triangle) and ADC-fMRI (n = 12, upper triangle). The gray matter ROIs and white matter ROIs are indicated by dark and light bars. Functional connectivity, defined as Fisher-transformed Pearson’s correlation coefficient, is indicated by the color bar.

**Supplementary Figure 12:**
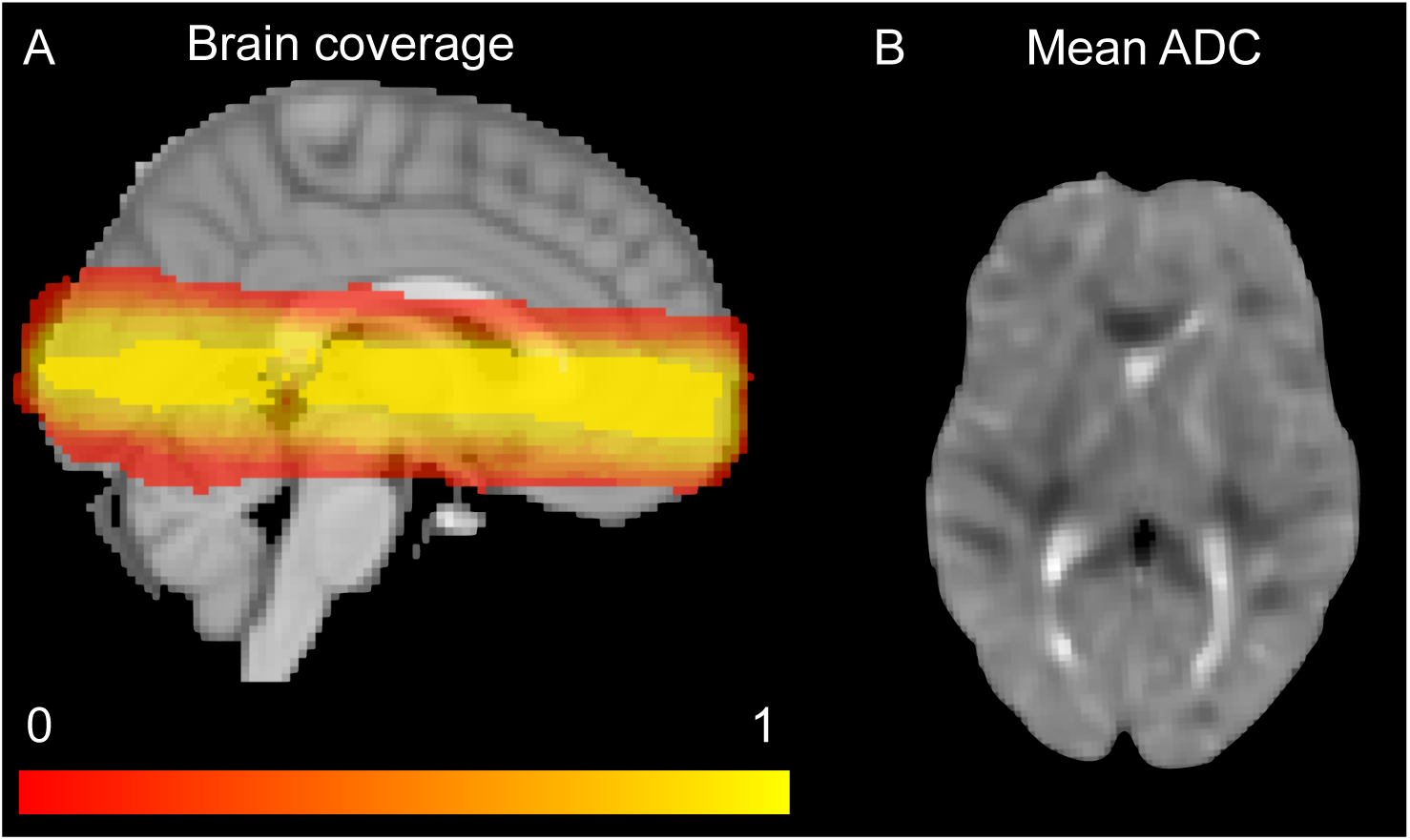
Illustration of partial brain coverage (A) and mean ADC (B). A) The brain coverage shows the average of each individual binary mask of brain coverage for ADC-fMRI at 3T (n = 12). The imaging slab was placed to encompass part of the DMN. The scale shows the percentage of overlap across subjects. The voxels that are common to all subjects are depicted in bright yellow. B) The mean ADC shows the temporal average of the ADC-fMRI of one subject at 3T.

**Supplementary Figure 13:**
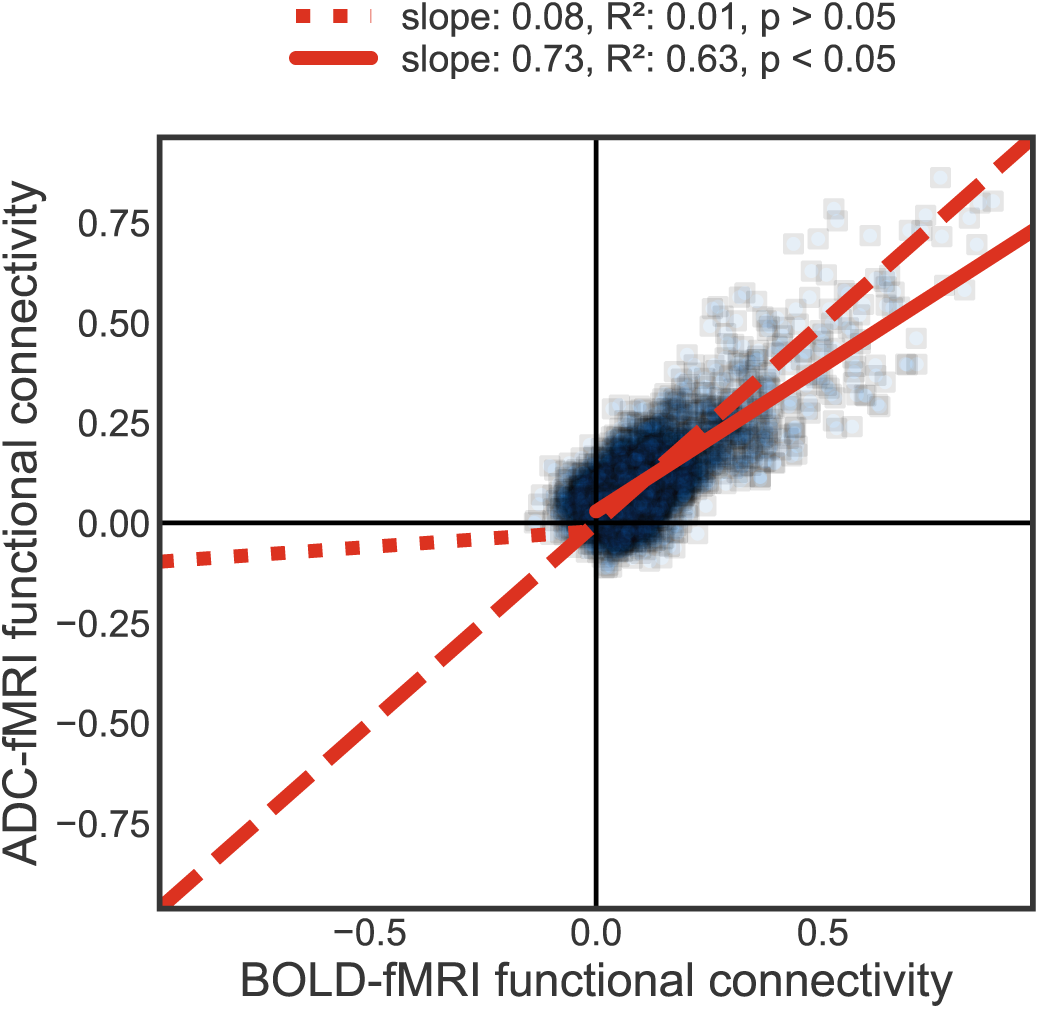
**No GSR, 3T: ADC-to-BOLD agreement of mean FC strength**, between corresponding edges. The solid line fits the positive correlations, while the dotted line fits the negative correlations. The dashed line represents a perfect agreement (identity line). Functional connectivity strength is defined as Fisher-transformed Pearson’s correlation coefficient. There is excellent agreement between ADC-fMRI and BOLD-fMRI positive FC, while negative FC is near absent.

**Supplementary Figure 14:**
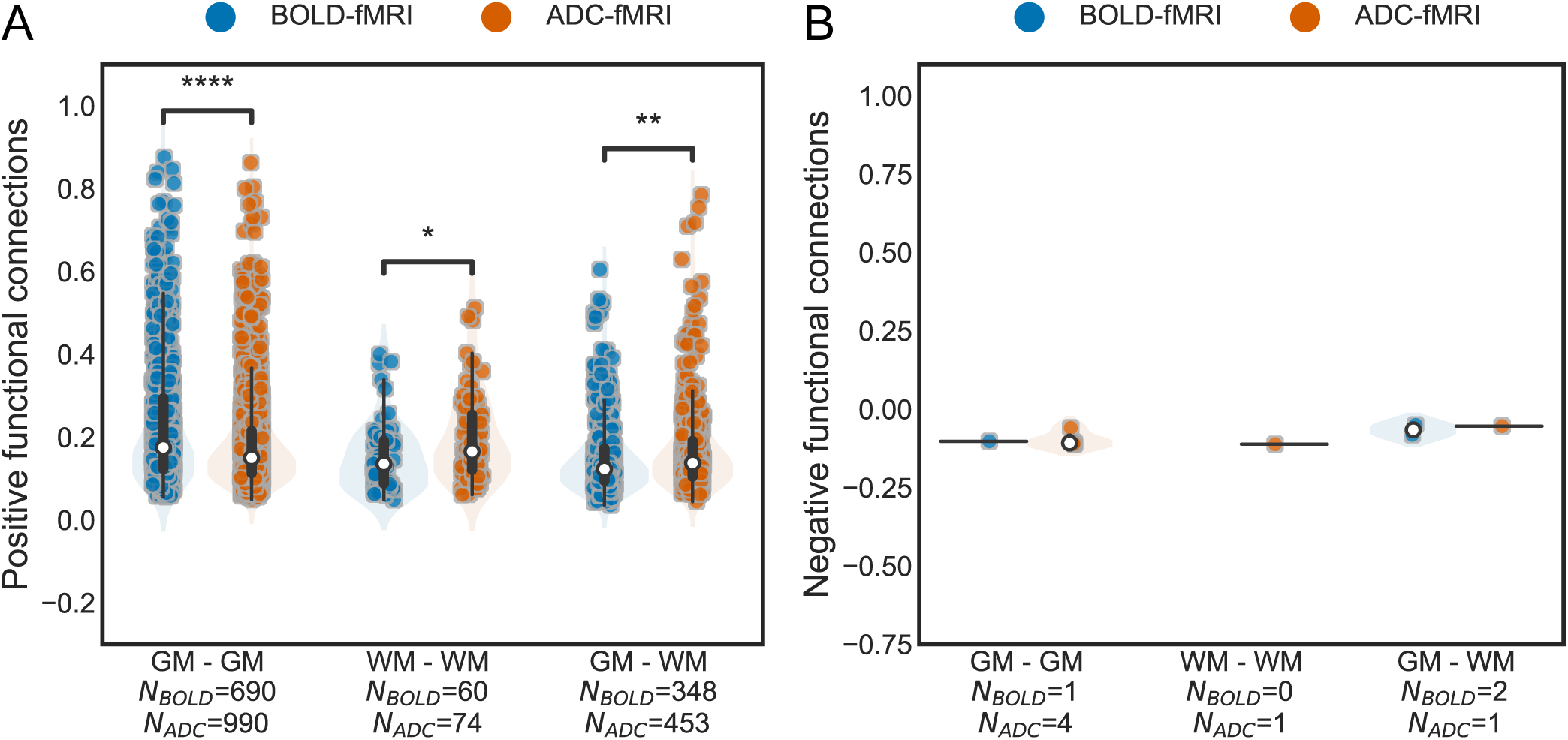
No GSR, 3T: Significant positive and negative group-averaged functional connections. The group-averaged correlations are split by tissue type (GM-GM, WM-WM, and GM-WM connectivity). Each dot corresponds to a significant edge of the FC, averaged across subjects. A)Positive correlations, B) negative correlations. The number of significant edges per functional contrast is indicated on the x-axis label. No negative correlations were found for BOLD-fMRI WM-WM. A box-and-whisker plot is shown within each violin plot, with a white dot for the median. Two-sided Mann-Whitney-Wilcoxon tests with Bonferroni correction are performed. * 0.01 *<* p ≤ 0.05; ** 0.001 *<* p ≤ 0.01; *** 0.0001 *<* p ≤ 0.001; **** p ≤ 0.0001.

**Supplementary Figure 15:**
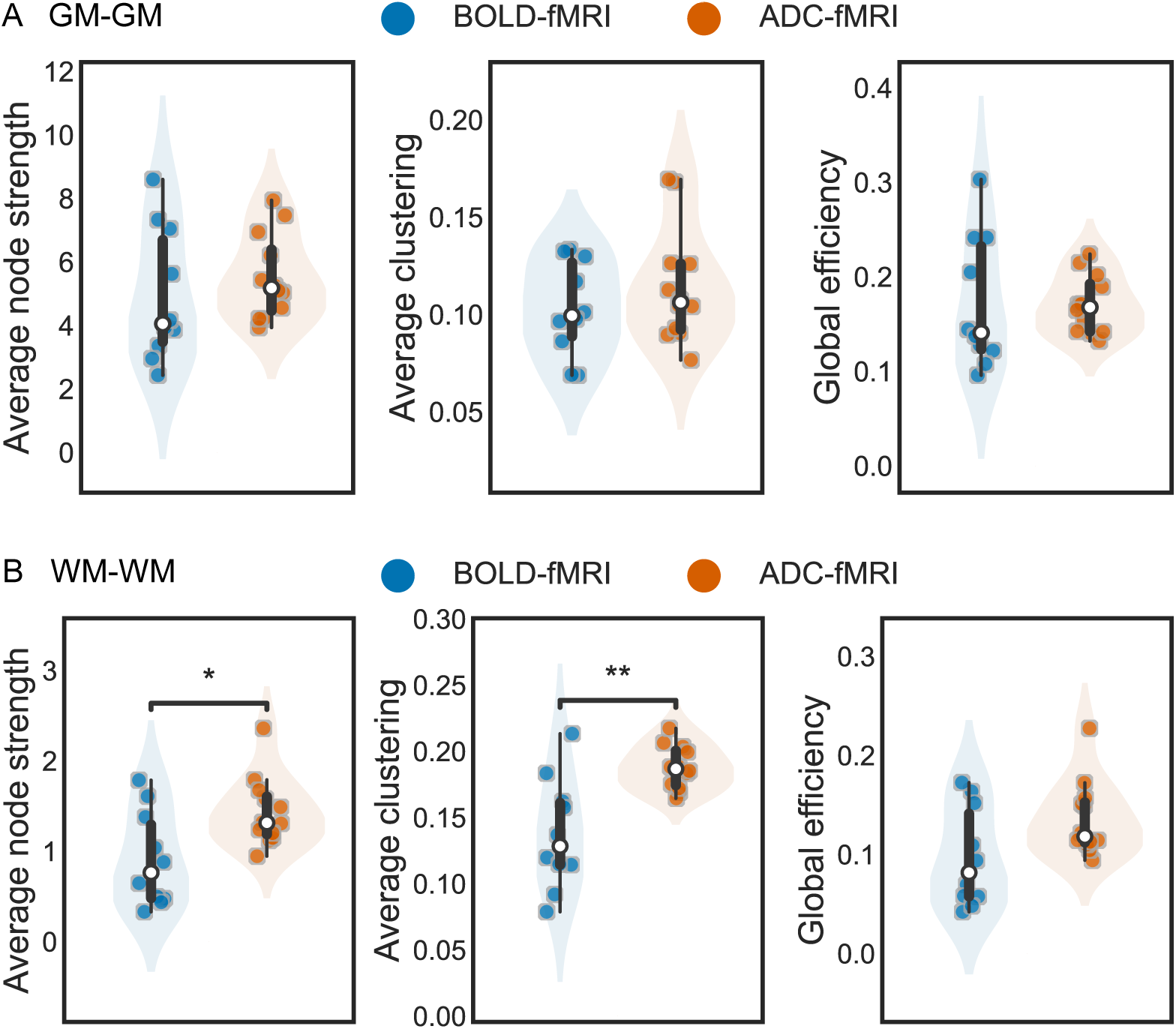
No GSR, 3T: Subject-wise weighted graph metrics. Graph metrics of BOLD-fMRI (n = 10) and ADC-fMRI (n = 12), based on A) GM-GM connectivity, or B) WM-WM connectivity. A box-and-whisker plot is shown within each violin plot, with a white dot for the median. Two-sided Mann-Whitney-Wilcoxon tests with Bonferroni correction were performed. * 0.01 *<* p ≤ 0.05; ** 0.001 *<* p ≤ 0.01; *** 0.0001 *<* p ≤ 0.001; **** p ≤ 0.0001.

**Supplementary Figure 16:**
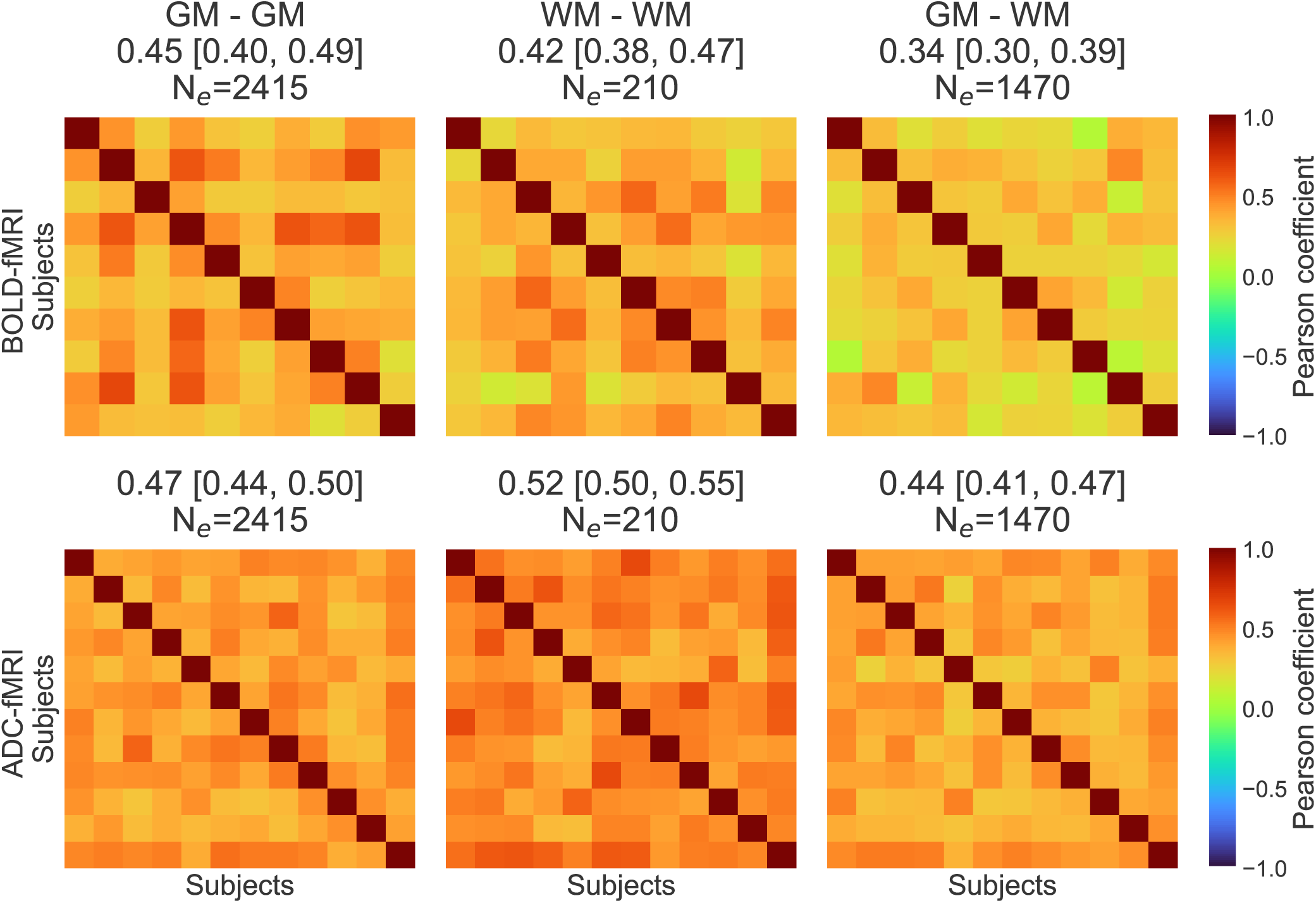
**All edges, 3T: Inter-subject FC similarity of all edges** for BOLD-fMRI and ADC-fMRI, calculated as the pairwise Pearson correlation of the vectorized FC matrices. Each similarity matrix has a size N*_subjects_* x N*_subjects_*, with N*_BOLD_*_−_*_fMRI_* = 10 and N*_ADC_*_−_*_fMRI_* = 12. The mean Pearson’s correlation coefficient of the similarity matrix and the 95% CI are depicted in the subtitles. N*_e_* corresponds to the number of FC edges on which the Pearson’s correlation coefficients are calculated. The GM-GM, WM-WM and GM-WM connectivity were considered separately.

**Supplementary Table 1:** SNR at 3T.

| Contrast | Whole brain | GM | WM |
| --- | --- | --- | --- |
| BOLD | $19.37 \pm 1.10$ | $20.16 \pm 1.15$ | $13.76 \pm 0.97$ |
| dfMRI 200 | $12.72 \pm 0.83$ | $13.18 \pm 0.87$ | $9.42 \pm 0.69$ |
| dfMRI 1000 | $7.39 \pm 0.45$ | $7.60 \pm 0.46$ | $5.93 \pm 0.51$ |

**Supplementary Table 2:**
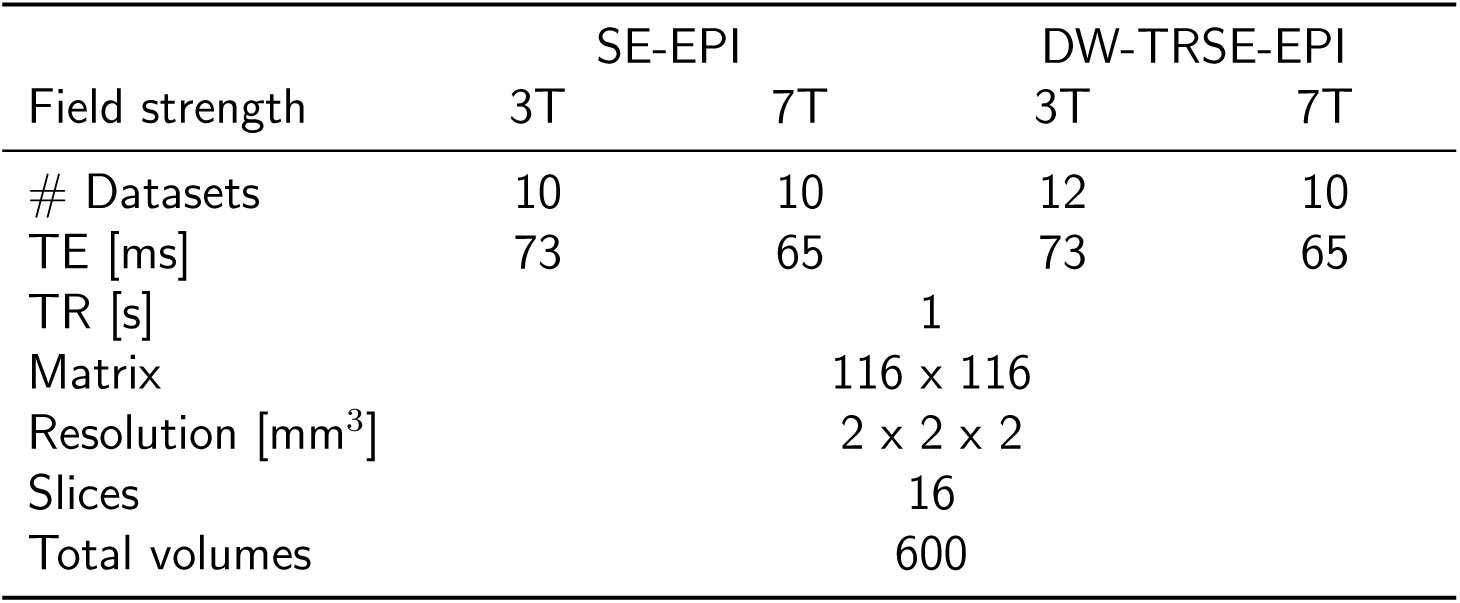
Acquisition parameters for SE-EPI and DW-TRSE-EPI. For DW-TRSE-EPI, interleaved volumes with b_1_ = 200 and b_2_ = 1000 s mm^−2^ were acquired. One b_1_ and one b_2_ volumes are needed to calculate one ADC volume, therefore doubling the sampling period from 1 s to 2 s for ADC-fMRI. Of the twenty-two subjects scanned at 3T and 7T, only 13 (BOLD 3T), 15 (ADC 3T), 16 (BOLD 7T) and 16 (ADC 7T) were acquired with the consistent sequence for rs. One subject dropped out of the study, one subject was excluded due to pronounced artifact resulting from a dental retainer and another because they fell asleep during the scanner. Six runs were discarded due to too many outlier time points (3 for ADC and 3 for BOLD, at 7T).

**Supplementary Table 3:**
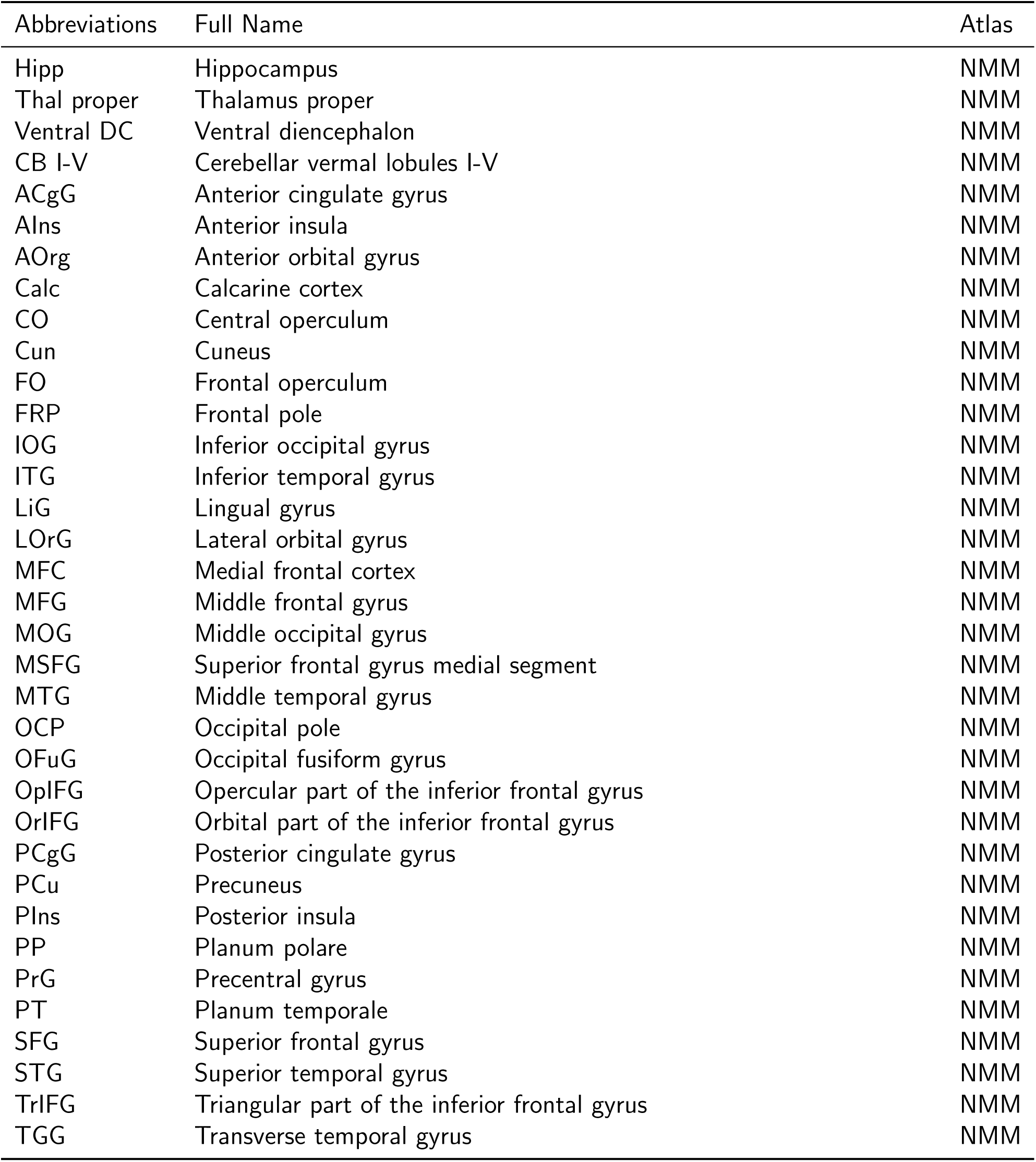
Abbreviations of the gray matter regions of interest. NMM = Neuromorphometrics. If R or L is added to the abbreviation, it corresponds to the right or left hemisphere, respectively.

**Supplementary Table 4:**
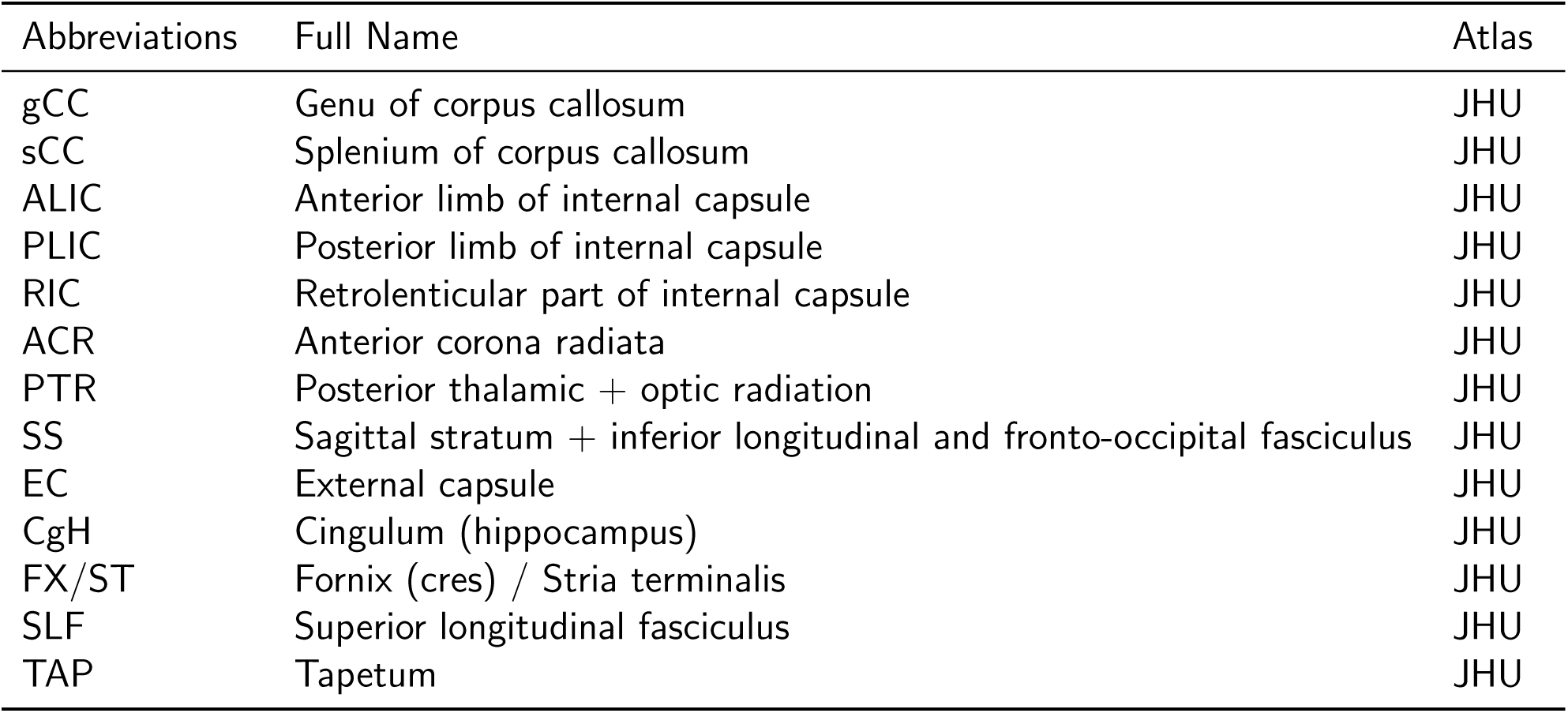
Abbreviations of the white matter regions of interest. JHU = John Hopkins University. If R or L is added to the abbreviation, it corresponds to the right or left hemisphere, respectively.

## Notes

### Competing Interest Statement

The authors have declared no competing interest.

### Summary of Updates

We have added a figure to clarify the methods (Figure 1), and a figure to visualize the functional connectivity of ADC-fMRI and BOLD-fMRI (Figure 2). We have updated the discussion with the effect of the SNR (Supplementary Table 1) and the effect of the higher sampling rate of BOLD-fMRI (Supplementary Figures 6-10) on the results.

## References

[1] B. Biswal, F. Z. Yetkin, V. M. Haughton, and J. S. Hyde. Functional connectivity in the motor cortex of resting human brain using echo-planar MRI. Magnetic Resonance in Medicine, 34(4): 537–541, October 1995. ISSN 0740-3194. doi: 10.1002/mrm.1910340409.

[2] Teddy Tjandra, Jonathan C. W. Brooks, Patricia Figueiredo, Richard Wise, Paul M. Matthews, and Irene Tracey. Quantitative assessment of the reproducibility of functional activation measured with BOLD and MR perfusion imaging: Implications for clinical trial design. NeuroImage, 27 (2):393–401, August 2005. ISSN 1053-8119. doi: 10.1016/j.neuroimage.2005.04.021. URL https://www.sciencedirect.com/science/article/pii/S1053811905002612.

[3] José M. Soares, Ricardo Magalhães, Pedro S. Moreira, Alexandre Sousa, Edward Ganz, Adriana Sampaio, Victor Alves, Paulo Marques, and Nuno Sousa. A Hitchhiker’s Guide to Functional Magnetic Resonance Imaging. Frontiers in Neuroscience, 10, 2016. ISSN 1662-453X. doi: 10.3389/fnins.2016.00515. URL https://www.frontiersin.org/articles/10.3389/fnins.2016.00515.

[4] Marcus E. Raichle, Ann Mary MacLeod, Abraham Z. Snyder, William J. Powers, Debra A. Gusnard, and Gordon L. Shulman. A default mode of brain function. Proceedings of the National Academy of Sciences, 98(2):676–682, January 2001. doi: 10.1073/pnas.98.2.676. URL https://www.pnas.org/doi/10.1073/pnas.98.2.676.

[5] Parinaz Babaeeghazvini, Laura M. Rueda-Delgado, Jolien Gooijers, Stephan P. Swinnen, and Andreas Daffertshofer. Brain Structural and Functional Connectivity: A Review of Combined Works of Diffusion Magnetic Resonance Imaging and Electro-Encephalography. Frontiers in Human Neuroscience, 15, 2021. ISSN 1662-5161. doi: 10.3389/fnhum.2021.721206. URL https://www.frontiersin.org/articles/10.3389/fnhum.2021.721206.

[6] H. Lv, Z. Wang, E. Tong, L.M. Williams, G. Zaharchuk, M. Zeineh, A.N. Goldstein-Piekarski, T.M. Ball, C. Liao, and M. Wintermark. Resting-State Functional MRI: Everything That Nonexperts Have Always Wanted to Know. AJNR: American Journal of Neuroradiology, 39(8):1390–1399, August 2018. ISSN 0195-6108. doi: 10.3174/ajnr.A5527. URL https://www.ncbi.nlm.nih.gov/pmc/articles/PMC6051935/.

[7] Martijn P. van den Heuvel and Hilleke E. Hulshoff Pol. Exploring the brain network: A review on resting-state fMRI functional connectivity. European Neuropsychopharmacology, 20(8):519–534 August 2010. ISSN 0924-977X. doi: 10.1016/j.euroneuro.2010.03.008. URL https://www.sciencedirect.com/science/article/pii/S0924977X10000684.

[8] Dongyang Zhang, James M. Johnston, Michael D. Fox, Eric C. Leuthardt, Robert L. Grubb, Michael R. Chicoine, Matthew D. Smyth, Abraham Z. Snyder, Marcus E. Raichle, and Joshua S. Shimony. Preoperative sensorimotor mapping in brain tumor patients using spontaneous fluctuations in neuronal activity imaged with functional magnetic resonance imaging: initial experience. Neurosurgery, 65(6 Suppl):226–236, December 2009. ISSN 1524-4040. doi: 10.1227/01.NEU.0000350868.95634.CA.

[9] Sara Stampacchia, Saina Asadi, Szymon Tomczyk, Federica Ribaldi, Max Scheffler, Karl-Olof Lövblad, Michela Pievani, Giovanni B. Frisoni, Valentina Garibotto, and Enrico Amico. Finger-printing of brain disease: Connectome identifiability in cognitive decline and neurodegeneration, February 2022. URL https://www.biorxiv.org/content/10.1101/2022.02.04.479112v1. Pages: 2022.02.04.479112 Section: New Results.

[10] Xi-Nian Zuo, Adriana Di Martino, Clare Kelly, Zarrar E. Shehzad, Dylan G. Gee, Donald F. Klein, F. Xavier Castellanos, Bharat B. Biswal, and Michael P. Milham. The oscillating brain: Complex and reliable. NeuroImage, 49(2):1432–1445 January 2010. ISSN 1053-8119. doi: 10.1016/j.neuroimage.2009.09.037. URL https://www.sciencedirect.com/science/article/pii/S1053811909010167.

[11] John C. Gore, Muwei Li, Yurui Gao, Tung-Lin Wu, Kurt G. Schilling, Yali Huang, Arabinda Mishra, Allen T. Newton, Baxter P. Rogers, Li Min Chen, Adam W. Anderson, and Zhaohua Ding. Functional MRI and resting state connectivity in white matter - a mini-review. Magnetic Resonance Imaging, 63:1–11 November 2019. ISSN 1873-5894. doi: 10.1016/j.mri.2019.07.017.

[12] Kathryn L. West, Mark D. Zuppichini, Monroe P. Turner, Dinesh K. Sivakolundu, Yuguang Zhao, Dema Abdelkarim, Jeffrey S. Spence, and Bart Rypma. BOLD hemodynamic response function changes significantly with healthy aging. NeuroImage, 188:198–207 March 2019. ISSN 1095-9572. doi: 10.1016/j.neuroimage.2018.12.012.

[13] Rebecca W. Pak, Darian H. Hadjiabadi, Janaka Senarathna, Shruti Agarwal, Nitish V. Thakor, Jay J. Pillai, and Arvind P. Pathak. Implications of neurovascular uncoupling in functional magnetic resonance imaging (fMRI) of brain tumors. Journal of Cerebral Blood Flow and Metabolism: Official Journal of the International Society of Cerebral Blood Flow and Metabolism, 37(11):3475–3487 November 2017. ISSN 1559-7016. doi: 10.1177/0271678X17707398.

[14] Christian Fynbo Christiansen. Risk of vascular disease in patients with multiple sclerosis: a review. Neurological Research, 34(8):746–753 October 2012. ISSN 1743-1328. doi: 10.1179/ 1743132812Y.0000000051.

[15] Toshihiko Aso, Shin-ichi Urayama, Hidenao Fukuyama, and Denis Le Bihan. Comparison of diffusion-weighted fMRI and BOLD fMRI responses in a verbal working memory task. NeuroImage, 67:25–32 February 2013. ISSN 1053-8119. doi: 10.1016/j.neuroimage.2012.11.005. URL https://www.sciencedirect.com/science/article/pii/S1053811912011019.

[16] Yoshifumi Abe, Norio Takata, Yuki Sakai, Hiro Taiyo Hamada, Yuichi Hiraoka, Tomomi Aida, Kohichi Tanaka, Denis Le Bihan, Kenji Doya, and Kenji F Tanaka. Diffusion functional MRI reveals global brain network functional abnormalities driven by targeted local activity in a neuropsychiatric disease mouse model. NeuroImage, 223:117318, December 2020. ISSN 1053-8119. doi: 10.1016/ j.neuroimage.2020.117318. URL https://www.sciencedirect.com/science/article/pii/ S1053811920308041.

[17] Clarisse I. Mark, Erin L. Mazerolle, and J. Jean Chen. Metabolic and vascular origins of the BOLD effect: Implications for imaging pathology and resting-state brain function. Journal of Magnetic Resonance Imaging, 42(2):231–246, 2015. ISSN 1522-2586. doi: 10.1002/jmri.24786. URL https://onlinelibrary.wiley.com/doi/abs/10.1002/jmri.24786.

[18] Lukas A. Grajauskas, Tory Frizzell, Xiaowei Song, and Ryan C. N. D’Arcy. White Matter fMRI Activation Cannot Be Treated as a Nuisance Regressor: Overcoming a Historical Blind Spot. Frontiers in Neuroscience, 13, 2019. ISSN 1662-453X. doi: DOI=10.3389/fnins.2019.01024. URL https://www.frontiersin.org/journals/neuroscience/articles/10.3389/fnins.2019.01024.

[19] Jodie R. Gawryluk, Erin L. Mazerolle, and Ryan C. N. D’Arcy. Does functional MRI detect activation in white matter? A review of emerging evidence, issues, and future directions. Frontiers in Neuroscience, 8, August 2014. ISSN 1662-453X. doi: 10.3389/fnins.2014.00239. URL https://www.frontiersin.org/journals/neuroscience/articles/10.3389/fnins.2014.00239/full.

[20] Muwei Li, Allen T. Newton, Adam W. Anderson, Zhaohua Ding, and John C. Gore. Characterization of the hemodynamic response function in white matter tracts for event-related fMRI. Nature Communications, 10(1):1140, March 2019. ISSN 2041-1723. doi: 10.1038/s41467-019-09076-2. URL https://www.nature.com/articles/s41467-019-09076-2.

[21] Yali Huang, Stephen K. Bailey, Peiguang Wang, Laurie E. Cutting, John C. Gore, and Zhaohua Ding. Voxel-wise Detection of Functional Networks in White Matter. NeuroImage, 183:544– 552 December 2018. ISSN 1053-8119. doi: 10.1016/j.neuroimage.2018.08.049. URL https://www.ncbi.nlm.nih.gov/pmc/articles/PMC6226032/.

[22] Zhaohua Ding, Yali Huang, Stephen K. Bailey, Yurui Gao, Laurie E. Cutting, Baxter P. Rogers, Allen T. Newton, and John C. Gore. Detection of synchronous brain activity in white matter tracts at rest and under functional loading. Proceedings of the National Academy of Sciences of the United States of America, 115(3):595–600 January 2018. ISSN 1091-6490. doi: 10.1073/pnas. 1711567115.

[23] Michael Peer, Mor Nitzan, Atira S. Bick, Netta Levin, and Shahar Arzy. Evidence for Functional Networks within the Human Brain’s White Matter. Journal of Neuroscience, 37(27):6394–6407 July 2017. ISSN 0270-6474, 1529-2401. doi: 10.1523/JNEUROSCI.3872-16.2017. URL https://www.jneurosci.org/content/37/27/6394.

[24] Jiao Li, Bharat B. Biswal, Pan Wang, Xujun Duan, Qian Cui, Huafu Chen, and Wei Liao. Exploring the functional connectome in white matter. Human Brain Mapping, 40(15):4331–4344, 2019. ISSN 1097-0193. doi: 10.1002/hbm.24705. URL https://onlinelibrary.wiley.com/doi/abs/10.1002/hbm.24705.

[25] Wiktor Olszowy, Yujian Diao, and Ileana O. Jelescu. Beyond BOLD: Evidence for diffusion fMRI contrast in the human brain distinct from neurovascular response. May 2021. doi: 10.1101/2021. 05.16.444253. Preprint at http://biorxiv.org/lookup/doi/10.1101/2021.05.16.444253.

[26] Anne Darquié, Jean-Baptiste Poline, Cyril Poupon, Hervé Saint-Jalmes, and Denis Le Bihan. Transient decrease in water diffusion observed in human occipital cortex during visual stimulation. Proceedings of the National Academy of Sciences, 98(16):9391–9395 July 2001. doi: 10.1073/ pnas.151125698. URL https://www.pnas.org/doi/abs/10.1073/pnas.151125698.

[27] Roland Bammer. Basic principles of diffusion-weighted imaging. European Journal of Radiology, 45(3):169–184 March 2003. ISSN 0720-048X. doi: 10.1016/S0720-048X(02)00303-0. URL https://www.sciencedirect.com/science/article/pii/S0720048X02003030.

[28] Daniel Nunes, Rita Gil, and Noam Shemesh. A rapid-onset diffusion functional MRI signal reflects neuromorphological coupling dynamics. NeuroImage, 231:117862, May 2021. ISSN 1053-8119. doi: 10.1016/j.neuroimage.2021.117862. URL https://www.sciencedirect.com/science/ article/pii/S1053811921001397.

[29] Tong Ling, Kevin C. Boyle, Valentina Zuckerman, Thomas Flores, Charu Ramakrishnan, Karl Deisseroth, and Daniel Palanker. High-speed interferometric imaging reveals dynamics of neuronal deformation during the action potential. Proceedings of the National Academy of Sciences, 117 (19):10278–10285 May 2020. doi: 10.1073/pnas.1920039117. URL https://www.pnas.org/doi/full/10.1073/pnas.1920039117.

[30] Junhwan Kwon, Sungho Lee, Yongjae Jo, and Myunghwan Choi. All-optical observation on activity-dependent nanoscale dynamics of myelinated axons. Neurophotonics, 10(1):015003, January 2023. ISSN 2329-423X. doi: 10.1117/1.NPh.10.1.015003. URL https://www.ncbi.nlm.nih.gov/ pmc/articles/PMC9868287/.

[31] Ronan Chéreau, G. Ezequiel Saraceno, Julie Angibaud, Daniel Cattaert, and U. Valentin Nagerl. Superresolution imaging reveals activity-dependent plasticity of axon morphology linked to changes in action potential conduction velocity. Proceedings of the National Academy of Sciences, 114(6): 1401–1406 February 2017. doi: 10.1073/pnas.1607541114. URL https://www.pnas.org/doi/ full/10.1073/pnas.1607541114.

[32] Ang Doma Sherpa, Fanrong Xiao, Neethu Joseph, Chiye Aoki, and Sabina Hrabetova. Activation of *β*-adrenergic receptors in rat visual cortex expands astrocytic processes and reduces extracellular space volume. *Synapse (New York*, N.Y.), 70(8):307–316 August 2016. ISSN 0887-4476. doi: 10.1002/syn.21908. URL https://www.ncbi.nlm.nih.gov/pmc/articles/PMC4909535/.

[33] Clement Debaker, Boucif Djemai, Luisa Ciobanu, Tomokazu Tsurugizawa, and Denis Le Bihan. Diffusion MRI reveals in vivo and non-invasively changes in astrocyte function induced by an aquaporin-4 inhibitor. PLOS ONE, 15(5):e0229702, May 2020. ISSN 1932-6203. doi: 10.1371/ journal.pone.0229702. URL https://journals.plos.org/plosone/article?id=10.1371/journal.pone.0229702. Publisher: Public Library of Science.

[34] Karla L. Miller, Daniel P. Bulte, Hannah Devlin, Matthew D. Robson, Richard G. Wise, Mark W. Woolrich, Peter Jezzard, and Timothy E. J. Behrens. Evidence for a vascular contribution to diffusion FMRI at high b value. Proceedings of the National Academy of Sciences, 104(52): 20967–20972 December 2007. doi: 10.1073/pnas.0707257105. URL https://www.pnas.org/doi/full/10.1073/pnas.0707257105.

[35] Ruiliang Bai, Craig V. Stewart, Dietmar Plenz, and Peter J. Basser. Assessing the sensitivity of diffusion MRI to detect neuronal activity directly. Proceedings of the National Academy of Sciences, 113(12):E1728–E1737 March 2016. doi: 10.1073/pnas.1519890113. URL https://www.pnas.org/doi/10.1073/pnas.1519890113.

[36] Yuji Komaki, Clement Debacker, Boucif Djemai, Luisa Ciobanu, Tomokazu Tsurugizawa, and Denis Le Bihan. Differential effects of aquaporin-4 channel inhibition on BOLD fMRI and diffusion fMRI responses in mouse visual cortex. PLoS ONE, 15(5):e0228759, May 2020. ISSN 1932-6203. doi: 10.1371/journal.pone.0228759. URL https://www.ncbi.nlm.nih.gov/pmc/articles/PMC7241787/.

[37] Tomokazu Tsurugizawa, Luisa Ciobanu, and Denis Le Bihan. Water diffusion in brain cortex closely tracks underlying neuronal activity. Proceedings of the National Academy of Sciences, 110(28): 11636–11641 July 2013. doi: 10.1073/pnas.1303178110. URL https://www.pnas.org/doi/10.1073/pnas.1303178110.

[38] André Pampel, Thies H. Jochimsen, and Harald E. Möller. BOLD background gradient contributions in diffusion-weighted fMRI–comparison of spin-echo and twice-refocused spin-echo sequences. NMR in biomedicine, 23(6):610–618 July 2010. ISSN 1099-1492. doi: 10.1002/nbm.1502.

[39] Denis Le Bihan, Shin-ichi Urayama, Toshihiko Aso, Takashi Hanakawa, and Hidenao Fukuyama. Direct and fast detection of neuronal activation in the human brain with diffusion MRI. Proceedings of the National Academy of Sciences of the United States of America, 103(21):8263–8268 May 2006. ISSN 0027-8424. doi: 10.1073/pnas.0600644103.

[40] Denis Le Bihan. What can we see with IVIM MRI? NeuroImage, 187:56–67 February 2019. ISSN 1053-8119. doi: 10.1016/j.neuroimage.2017.12.062. URL https://www.sciencedirect.com/science/article/pii/S1053811917310868.

[41] Mami Iima and Denis Le Bihan. Clinical Intravoxel Incoherent Motion and Diffusion MR Imaging: Past, Present, and Future. Radiology, 278(1):13–32 January 2016. ISSN 1527-1315. doi: 10.1148/radiol.2015150244.

[42] Yoshifumi Abe, Tomokazu Tsurugizawa, and Denis Le Bihan. Water diffusion closely reveals neural activity status in rat brain loci affected by anesthesia. PLoS Biology, 15(4):e2001494, April 2017. ISSN 1544-9173. doi: 10.1371/journal.pbio.2001494. URL https://www.ncbi.nlm.nih.gov/pmc/articles/PMC5390968/.

[43] William M. Spees, Tsen-Hsuan Lin, and Sheng-Kwei Song. White-matter diffusion fMRI of mouse optic nerve. NeuroImage, 65:209–215 January 2013. ISSN 1053-8119. doi: 10.1016/j.neuroimage.2012.10.021. URL https://www.sciencedirect.com/science/article/pii/S1053811912010245.

[44] Michael D. Fox, Abraham Z. Snyder, Justin L. Vincent, Maurizio Corbetta, David C. Van Essen, and Marcus E. Raichle. The human brain is intrinsically organized into dynamic, anticorrelated functional networks. Proceedings of the National Academy of Sciences, 102(27):9673–9678 July 2005. doi: 10.1073/pnas.0504136102. URL https://www.pnas.org/doi/full/10.1073/pnas.0504136102.

[45] Marta Bianciardi, Masaki Fukunaga, Peter van Gelderen, Jacco A de Zwart, and Jeff H Duyn. Negative BOLD-fMRI signals in large cerebral veins. Journal of Cerebral Blood Flow & Metabolism, 31(2):401–412 February 2011. ISSN 0271-678X. doi: 10.1038/jcbfm.2010.164. URL https://www.ncbi.nlm.nih.gov/pmc/articles/PMC3049531/.

[46] Parker G. Allan, Robert G. Briggs, Andrew K. Conner, Christen M. O’Neal, Phillip A. Bonney, Brian D. Maxwell, Cordell M. Baker, Joshua D. Burks, Goksel Sali, Chad A. Glenn, and Michael E. Sughrue. Parcellation-based tractographic modeling of the dorsal attention network. Brain and Behavior, 9(10):e01365, 2019. ISSN 2162-3279. doi: 10.1002/brb3.1365. URL https://onlinelibrary.wiley.com/doi/abs/10.1002/brb3.1365.

[47] Simone Vossel, Joy J. Geng, and Gereon R. Fink. Dorsal and Ventral Attention Systems: Distinct Neural Circuits but Collaborative Roles. The Neuroscientist, 20(2):150–159 April 2014. ISSN 1073-8584. doi: 10.1177/1073858413494269. URL 10.1177/1073858413494269.

[48] Pedro Nascimento Alves, Stephanie J. Forkel, Maurizio Corbetta, and Michel Thiebaut de Schotten. The subcortical and neurochemical organization of the ventral and dorsal attention networks. Communications Biology, 5(1):1–14 December 2022. ISSN 2399-3642. doi: 10.1038/s42003-022-04281-0. URL https://www.nature.com/articles/s42003-022-04281-0.

[49] Xiaoxiao Xu, Hong Yuan, and Xu Lei. Activation and Connectivity within the Default Mode Network Contribute Independently to Future-Oriented Thought. Scientific Reports, 6(1):21001, February 2016. ISSN 2045-2322. doi: 10.1038/srep21001. URL https://www.nature.com/ articles/srep21001.

[50] Weifang Cao, Cheng Luo, Bin Zhu, Dan Zhang, Li Dong, Jinnan Gong, Diankun Gong, Hui He, Shipeng Tu, Wenjie Yin, Jianfu Li, Huafu Chen, and Dezhong Yao. Resting-state functional connectivity in anterior cingulate cortex in normal aging. Frontiers in Aging Neuroscience, 6, 2014. ISSN 1663-4365. doi: 10.3389/fnagi.2014.00280. URL https://www.frontiersin.org/ articles/10.3389/fnagi.2014.00280.

[51] Michael D. Fox, Dongyang Zhang, Abraham Z. Snyder, and Marcus E. Raichle. The Global Signal and Observed Anticorrelated Resting State Brain Networks. Journal of Neurophysiology, 101(6):3270–3283 June 2009. ISSN 0022-3077. doi: 10.1152/jn.90777.2008. URL https://journals.physiology.org/doi/full/10.1152/jn.90777.2008.

[52] Andreas Weissenbacher, Christian Kasess, Florian Gerstl, Rupert Lanzenberger, Ewald Moser, and Christian Windischberger. Correlations and anticorrelations in resting-state functional connectivity MRI: A quantitative comparison of preprocessing strategies. NeuroImage, 47(4):1408– 1416 October 2009. ISSN 1053-8119. doi: 10.1016/j.neuroimage.2009.05.005. URL https://www.sciencedirect.com/science/article/pii/S105381190900487X.

[53] Kevin Murphy, Rasmus M. Birn, Daniel A. Handwerker, Tyler B. Jones, and Peter A. Bandettini. The impact of global signal regression on resting state correlations: Are anti-correlated networks introduced? NeuroImage, 44(3):893–905 February 2009. ISSN 1053-8119. doi: 10. 1016/j.neuroimage.2008.09.036. URL https://www.sciencedirect.com/science/article/ pii/S1053811908010264.

[54] Kevin Murphy and Michael D. Fox. Towards a consensus regarding global signal regression for resting state functional connectivity MRI. NeuroImage, 154:169–173 July 2017. ISSN 1053-8119. doi: 10.1016/j.neuroimage.2016.11.052. URL https://www.sciencedirect.com/science/ article/pii/S1053811916306711.

[55] Jingwei Li, Ru Kong, Raphaël Liégeois, Csaba Orban, Yanrui Tan, Nanbo Sun, Avram J. Holmes, Mert R. Sabuncu, Tian Ge, and B. T. Thomas Yeo. Global signal regression strengthens association between resting-state functional connectivity and behavior. NeuroImage, 196: 126–141 August 2019. ISSN 1053-8119. doi: 10.1016/j.neuroimage.2019.04.016. URL https://www.sciencedirect.com/science/article/pii/S1053811919303027.

[56] Xiaoqian J. Chai, Alfonso Nieto Castañón, Dost Öngür, and Susan Whitfield-Gabrieli. Anticorrelations in resting state networks without global signal regression. NeuroImage, 59(2): 1420–1428 January 2012. ISSN 1053-8119. doi: 10.1016/j.neuroimage.2011.08.048. URL https://www.sciencedirect.com/science/article/pii/S1053811911009657.

[57] Jingyuan E. Chen, Laura D. Lewis, Catie Chang, Qiyuan Tian, Nina E. Fultz, Ned A. Ohringer, Bruce R. Rosen, and Jonathan R. Polimeni. Resting-state “physiological networks”. NeuroImage, 213:116707, June 2020. ISSN 1053-8119. doi: 10.1016/j.neuroimage.2020.116707. URL https://www.sciencedirect.com/science/article/pii/S1053811920301944.

[58] Guangyu Chen, Gang Chen, Chunming Xie, and Shi-Jiang Li. Negative Functional Connectivity and Its Dependence on the Shortest Path Length of Positive Network in the Resting-State Human Brain. Brain Connectivity, 1(3):195–206 September 2011. ISSN 2158-0014. doi: 10.1089/brain. 2011.0025. URL https://www.ncbi.nlm.nih.gov/pmc/articles/PMC3572722/.

[59] Setsu Wakana, Hangyi Jiang, Lidia M. Nagae-Poetscher, Peter C. M. van Zijl, and Susumu Mori. Fiber Tract–based Atlas of Human White Matter Anatomy. Radiology, 230(1):77–87 January 2004. ISSN 0033-8419. doi: 10.1148/radiol.2301021640. URL https://pubs.rsna.org/doi/ 10.1148/radiol.2301021640.

[60] I. Fatih Karababa, Huseyin Bayazıt, Nihat Kılıçaslan, Mustafa Celik, Hasan Cece, Ekrem Karakas, and Salih Selek. Microstructural Changes of Anterior Corona Radiata in Bipolar Depression. Psychiatry Investigation, 12(3):367–371 July 2015. ISSN 1738-3684. doi: 10.4306/pi.2015.12.3.367. URL https://www.ncbi.nlm.nih.gov/pmc/articles/PMC4504920/.

[61] Piero Chiacchiaretta and Antonio Ferretti. Resting State BOLD Functional Connectivity at 3T: Spin Echo versus Gradient Echo EPI. PLOS ONE, 10(3):e0120398, March 2015. ISSN 1932-6203. doi: 10.1371/journal.pone.0120398. URL https://journals.plos.org/plosone/article?id=10.1371/journal.pone.0120398.

[62] Emily S. Finn, Xilin Shen, Dustin Scheinost, Monica D. Rosenberg, Jessica Huang, Marvin M. Chun, Xenophon Papademetris, and R. Todd Constable. Functional connectome fingerprinting: identifying individuals using patterns of brain connectivity. Nature Neuroscience, 18(11):1664– 1671 November 2015. ISSN 1546-1726. doi: 10.1038/nn.4135. URL https://www.nature.com/articles/nn.4135.

[63] Tao Jin, Fuqiang Zhao, and Seong-Gi Kim. Sources of functional apparent diffusion coefficient changes investigated by diffusion-weighted spin-echo fMRI. Magnetic Resonance in Medicine, 56(6):1283–1292, 2006. ISSN 1522-2594. doi: 10.1002/mrm.21074. URL https://onlinelibrary.wiley.com/doi/abs/10.1002/mrm.21074.

[64] Tao Jin and Seong-Gi Kim. Functional changes of apparent diffusion coefficient during visual stimulation investigated by diffusion-weighted gradient-echo fMRI. NeuroImage, 41(3): 801–812 July 2008. ISSN 1053-8119. doi: 10.1016/j.neuroimage.2008.03.014. URL https://www.sciencedirect.com/science/article/pii/S1053811908002358.

[65] Ileana Ozana Jelescu, Luisa Ciobanu, Françoise Geffroy, Pierre Marquet, and Denis Le Bihan. Effects of hypotonic stress and ouabain on the apparent diffusion coefficient of water at cellular and tissue levels in Aplysia. NMR in Biomedicine, 27(3):280–290, 2014. ISSN 1099-1492. doi: 10.1002/nbm.3061. URL https://onlinelibrary.wiley.com/doi/abs/10.1002/nbm.3061.

[66] Joseph S. Gati, Ravi S. Menon, Kămil Uğurbil, and Brian K. Rutt. Experimental determination of the BOLD field strength dependence in vessels and tissue. Magnetic Resonance in Medicine, 38(2):296–302, 1997. ISSN 1522-2594. doi: 10.1002/mrm.1910380220. URL https://onlinelibrary.wiley.com/doi/abs/10.1002/mrm.1910380220. eprint: https://onlinelibrary.wiley.com/doi/pdf/10.1002/mrm.1910380220.

[67] Markus Barth and Benedikt A. Poser. Advances in High-Field BOLD fMRI. Materials, 4(11): 1941–1955 November 2011. ISSN 1996-1944. doi: 10.3390/ma4111941. URL https://www.mdpi.com/1996-1944/4/11/1941.

[68] Torben E. Lund, Kristoffer H. Madsen, Karam Sidaros, Wen-Lin Luo, and Thomas E. Nichols. Non-white noise in fMRI: Does modelling have an impact? NeuroImage, 29(1):54–66 January 2006. ISSN 1053-8119. doi: 10.1016/j.neuroimage.2005.07.005. URL https://www.sciencedirect. com/science/article/pii/S105381190500501X.

[69] Arthur P. C. Spencer, Jasmine Nguyen-Duc, Ines de Riedmatten, Filip Szczepankiewicz, and Ileana O. Jelescu. Mapping grey and white matter activity in the human brain with isotropic ADC-fMRI. October 2024. doi: 10.1101/2024.10.01.615823. Preprint at https://www.biorxiv.org/content/10.1101/2024.10.01.615823v1.

[70] Kawin Setsompop, Qiuyun Fan, Jason Stockmann, Berkin Bilgic, Susie Huang, Stephen F. Cauley, Aapo Nummenmaa, Fuyixue Wang, Yogesh Rathi, Thomas Witzel, and Lawrence L. Wald. High-resolution in vivo diffusion imaging of the human brain with Generalized SLIce Dithered Enhanced Resolution Simultaneous MultiSlice (gSlider-SMS). Magnetic resonance in medicine, 79 (1):141, January 2018. doi: 10.1002/mrm.26653. URL https://www.ncbi.nlm.nih.gov/pmc/articles/PMC5585027/.

[71] Tsen-Hsuan Lin, William M. Spees, Chia-Wen Chiang, Kathryn Trinkaus, Anne H. Cross, and Sheng-Kwei Song. Diffusion fMRI detects white-matter dysfunction in mice with acute optic neuritis. Neurobiology of Disease, 67:1–8 July 2014. ISSN 0969-9961. doi: 10.1016/j.nbd.2014.02.007. URL https://www.sciencedirect.com/science/article/pii/S0969996114000497.

[72] William M. Spees, Tsen-Hsuan Lin, Peng Sun, Chunyu Song, Ajit George, Sam E. Gary, Hsin-Chieh Yang, and Sheng-Kwei Song. MRI-based assessment of function and dysfunction in myelinated axons. Proceedings of the National Academy of Sciences, 115(43):E10225–E10234 October 2018. doi: 10.1073/pnas.1801788115. URL https://www.pnas.org/doi/abs/10.1073/pnas. 1801788115.

[73] Gang Zheng and William S. Price. Suppression of background gradients in (B0 gradient-based) NMR diffusion experiments. Concepts in Magnetic Resonance Part A, 30A(5):261–277, 2007. ISSN 1552-5023. doi: 10.1002/cmr.a.20092. URL https://onlinelibrary.wiley.com/doi/abs/10.1002/cmr.a.20092.

[74] Jerrold L. Boxerman, Leena M. Hamberg, Bruce R. Rosen, and Robert M. Weisskoff. Mr contrast due to intravascular magnetic susceptibility perturbations. Magnetic Resonance in Medicine, 34(4):555–566, 1995. ISSN 1522-2594. doi: 10.1002/mrm.1910340412. URL https://onlinelibrary.wiley.com/doi/abs/10.1002/mrm.1910340412.

[75] David G. Norris. Spin-echo fMRI: The poor relation? NeuroImage, 62(2):1109–1115 August 2012. ISSN 1053-8119. doi: 10.1016/j.neuroimage.2012.01.003. URL https://www.sciencedirect.com/science/article/pii/S1053811912000067.

[76] Bernd André Jung and Matthias Weigel. Spin echo magnetic resonance imaging. Journal of Magnetic Resonance Imaging, 37(4):805–817, 2013. ISSN 1522-2586. doi: 10.1002/jmri.24068. URL https://onlinelibrary.wiley.com/doi/abs/10.1002/jmri.24068.

[77] Birn RM and Bandettini PA. The effect of t2’ changes on spin-echo epi derived brain activation maps. In Proc. Intl. Soc. Mag. Reson. Med., Hawaii, 2002. URL https://cds.ismrm.org/ ismrm-2002/PDF5/1324.PDF. p. 1324.

[78] Keilholz SD, Silva AC, Duyn JH, and Koretsky AP. The contribution of t2* to spin-echo epi: implications for high-field fmri studies. In *Proc. Intl. Soc. Mag. Reson. Med.*, Miami, 2005. URL https://cds.ismrm.org/protected/05MProceedings/PDFfiles/00032.pdf. p. 32.

[79] Peter J. Koopmans, Rasim Boyacioğlu, Markus Barth, and David G. Norris. Whole brain, high resolution spin-echo resting state fMRI using PINS multiplexing at 7T. NeuroImage, 62(3):1939– 1946 September 2012. ISSN 1053-8119. doi: 10.1016/j.neuroimage.2012.05.080. URL https://www.sciencedirect.com/science/article/pii/S105381191200568X.

[80] Mark A. Griswold, Peter M. Jakob, Robin M. Heidemann, Mathias Nittka, Vladimir Jellus, Jianmin Wang, Berthold Kiefer, and Axel Haase. Generalized autocalibrating partially parallel acquisitions (GRAPPA). Magnetic Resonance in Medicine, 47(6):1202–1210, 2002. ISSN 1522-2594. doi: 10.1002/mrm.10171. URL https://onlinelibrary.wiley.com/doi/abs/10.1002/mrm.10171.

[81] K. Setsompop, J. Cohen-Adad, B. A. Gagoski, T. Raij, A. Yendiki, B. Keil, V. J. Wedeen, and L. L. Wald. Improving diffusion MRI using simultaneous multi-slice echo planar imaging. NeuroImage, 63(1):569–580 October 2012. ISSN 1053-8119. doi: 10.1016/j.neuroimage.2012.06.033. URL https://www.sciencedirect.com/science/article/pii/S1053811912006477.

[82] Kawin Setsompop, Borjan A. Gagoski, Jonathan R. Polimeni, Thomas Witzel, Van J. Wedeen, and Lawrence L. Wald. Blipped-controlled aliasing in parallel imaging for simultaneous multislice echo planar imaging with reduced g-factor penalty. Magnetic Resonance in Medicine, 67(5):1210–1224, 2012. ISSN 1522-2594. doi: 10.1002/mrm.23097. URL https://onlinelibrary.wiley.com/doi/abs/10.1002/mrm.23097.

[83] Kamil Uğurbil, Junqian Xu, Edward J. Auerbach, Steen Moeller, An T. Vu, Julio M. Duarte-Carvajalino, Christophe Lenglet, Xiaoping Wu, Sebastian Schmitter, Pierre Francois Van de Moortele, John Strupp, Guillermo Sapiro, Federico De Martino, Dingxin Wang, Noam Harel, Michael Garwood, Liyong Chen, David A. Feinberg, Stephen M. Smith, Karla L. Miller, Stamatios N. Sotiropoulos, Saad Jbabdi, Jesper L. R. Andersson, Timothy E. J. Behrens, Matthew F. Glasser, David C. Van Essen, Essa Yacoub, and WU-Minn HCP Consortium. Pushing spatial and temporal resolution for functional and diffusion MRI in the Human Connectome Project. NeuroImage, 80:80–104 October 2013. ISSN 1095-9572. doi: 10.1016/j.neuroimage.2013.05.012.

[84] Jelle Veraart, Dmitry S. Novikov, Daan Christiaens, Benjamin Ades-aron, Jan Sijbers, and Els Fieremans. Denoising of diffusion MRI using random matrix theory. NeuroImage, 142:394–406 November 2016. ISSN 1053-8119. doi: 10.1016/j.neuroimage.2016.08.016. URL https://www.ncbi.nlm.nih.gov/pmc/articles/PMC5159209/.

[85] Yujian Diao, Ting Yin, Rolf Gruetter, and Ileana O. Jelescu. PIRACY: An Optimized Pipeline for Functional Connectivity Analysis in the Rat Brain. Frontiers in Neuroscience, 15, 2021. ISSN 1662-453X. doi: 10.3389/fnins.2021.602170. URL https://www.frontiersin.org/articles/10.3389/fnins.2021.602170.

[86] Benjamin Ades-Aron, Gregory Lemberskiy, Jelle Veraart, John Golfinos, Els Fieremans, Dmitry S. Novikov, and Timothy Shepherd. Improved Task-based Functional MRI Language Mapping in Patients with Brain Tumors through Marchenko-Pastur Principal Component Analysis Denoising. Radiology, 298(2):365–373 February 2021. ISSN 0033-8419. doi: 10.1148/radiol.2020200822. URL https://pubs.rsna.org/doi/full/10.1148/radiol.2020200822.

[87] Elias Kellner, Bibek Dhital, Valerij G. Kiselev, and Marco Reisert. Gibbs-ringing artifact removal based on local subvoxel-shifts. Magnetic Resonance in Medicine, 76(5):1574–1581 November 2016. ISSN 1522-2594. doi: 10.1002/mrm.26054.

[88] Jesper L. R. Andersson, Stefan Skare, and John Ashburner. How to correct susceptibility distortions in spin-echo echo-planar images: application to diffusion tensor imaging. NeuroImage, 20(2):870– 888 October 2003. ISSN 1053-8119. doi: 10.1016/S1053-8119(03)00336-7.

[89] Fidel Alfaro-Almagro, Mark Jenkinson, Neal K. Bangerter, Jesper L. R. Andersson, Ludovica Griffanti, Gwenaëlle Douaud, Stamatios N. Sotiropoulos, Saad Jbabdi, Moises Hernandez-Fernandez, Emmanuel Vallee, Diego Vidaurre, Matthew Webster, Paul McCarthy, Christopher Rorden, Alessandro Daducci, Daniel C. Alexander, Hui Zhang, Iulius Dragonu, Paul M. Matthews, Karla L. Miller, and Stephen M. Smith. Image processing and Quality Control for the first 10,000 brain imaging datasets from UK Biobank. NeuroImage, 166:400–424, February 2018. ISSN 1095-9572. doi: 10.1016/j.neuroimage.2017.10.034.

[90] William D. Penny, Karl J. Friston, John T. Ashburner, Stefan J. Kiebel, and Thomas E. Nichols. Statistical Parametric Mapping: The Analysis of Functional Brain Images. Elsevier, April 2011. ISBN 978-0-08-046650-7.

[91] Brian B Avants, Nick Tustison, Gang Song, et al. Advanced normalization tools (ants). Insight j, 2(365):1–35, 2009.

[92] C.F. Beckmann and S.M. Smith. Probabilistic independent component analysis for functional magnetic resonance imaging. IEEE Transactions on Medical Imaging, 23(2):137–152 February 2004. ISSN 1558-254X. doi: 10.1109/TMI.2003.822821. URL https://ieeexplore.ieee.org/document/1263605.

[93] Ludovica Griffanti, Gwenaëlle Douaud, Janine Bijsterbosch, Stefania Evangelisti, Fidel Alfaro-Almagro, Matthew F. Glasser, Eugene P. Duff, Sean Fitzgibbon, Robert Westphal, Davide Carone, Christian F. Beckmann, and Stephen M. Smith. Hand classification of fMRI ICA noise components. Neuroimage, 154:188–205 July 2017. ISSN 1053-8119. doi: 10.1016/j.neuroimage.2016.12.036. URL https://www.ncbi.nlm.nih.gov/pmc/articles/PMC5489418/.

[94] Marieke L. Schölvinck, Alexander Maier, Frank Q. Ye, Jeff H. Duyn, and David A. Leopold. Neural basis of global resting-state fMRI activity. Proceedings of the National Academy of Sciences, 107 (22):10238–10243 June 2010. doi: 10.1073/pnas.0913110107. URL https://www.pnas.org/ doi/full/10.1073/pnas.0913110107.

[95] Yoav Benjamini and Yosef Hochberg. Controlling the False Discovery Rate: A Practical and Powerful Approach to Multiple Testing. Journal of the Royal Statistical Society. Series B (Methodological), 57(1):289–300, 1995. ISSN 0035-9246. URL https://www.jstor.org/stable/2346101.

[96] Mikail Rubinov and Olaf Sporns. Complex network measures of brain connectivity: Uses and interpretations. NeuroImage, 52(3):1059–1069 September 2010. ISSN 1053-8119. doi: 10.1016/j.neuroimage.2009.10.003. URL https://www.sciencedirect.com/science/article/pii/S105381190901074X.

[97] Duncan J. Watts and Steven H. Strogatz. Collective dynamics of ‘small-world’ networks. Nature, 393(6684):440–442, June 1998. ISSN 1476-4687. doi: 10.1038/30918. URL https://www.nature.com/articles/30918.

[98] Farzad V. Farahani, Waldemar Karwowski, and Nichole R. Lighthall. Application of Graph Theory for Identifying Connectivity Patterns in Human Brain Networks: A Systematic Review. Frontiers in Neuroscience, 13, 2019. ISSN 1662-453X. doi: 10.3389/fnins.2019.00585. URL https://www.frontiersin.org/journals/neuroscience/articles/10.3389/fnins.2019.00585.

[99] Olaf Sporns, Dante R. Chialvo, Marcus Kaiser, and Claus C. Hilgetag. Organization, development and function of complex brain networks. Trends in Cognitive Sciences, 8(9):418–425 September 2004. ISSN 1364-6613. doi: 10.1016/j.tics.2004.07.008. URL https://www.sciencedirect. com/science/article/pii/S1364661304001901.

[100] Wiktor Olszowy and Ileana Jelescu. diffusion fmri [dataset]. OpenNeuro, 2021. doi: 10.18112/openneuro.ds003676.v1.0.1. URL https://openneuro.org/datasets/ds003676/versions/1.0.1.

[101] In’es de Riedmatten, Arthur Spencer, Wiktor Olszowy, and Ileana O Jelescu. Apparent diffusion coefficient fmri shines light on white matter resting-state connectivity compared to bold [dataset]. figshare, 2025. doi: 10.6084/m9.figshare.28435325.v2.

[102] Wiktor Olszowy. Dfmri (v1.0.1) [computer software]. Github, February 2025. doi: 10.5281/zenodo.14935326. URL https://github.com/Mic-map/DfMRI/.

[103] In’es de Riedmatten. Adc rsfmri (v1.0.0) [computer software]. Github, February 2025. doi: 10.5281/zenodo.14920202. URL https://github.com/Mic-map/ADC_rsfMRI/.

[104] Martin Krzywinski, Jacqueline Schein, Inanç Birol, Joseph Connors, Randy Gascoyne, Doug Horsman, Steven J. Jones, and Marco A. Marra. Circos: an information aesthetic for comparative genomics. Genome Research, 19(9):1639–1645 September 2009. ISSN 1549-5469. doi: 10.1101/gr.092759.109.

